# Rapid acquisition of HIV-1 neutralization breadth in a rhesus V2 apex germline antibody mouse model after a single bolus immunization

**DOI:** 10.1101/2025.06.12.659380

**Authors:** Amrit Raj Ghosh, Rumi Habib, Nitesh Mishra, Ryan S. Roark, Madhav Akauliya, Ali A. Albowaidey, Joel D. Allen, Khaled Amereh, Gabriel Avillion, Maria Bottermann, Bo Liang, Namit Chaudhary, Sean Callaghan, Jonathan Dye, Xuduo Li, Jordan R. Ellis-Pugh, Rohan Roy Chowdhury, Nicole E. James, Xiaotie Liu, Laura Maiorino, Paula Maldonado, Rebecca Nedellec, Prabhgun Oberoi, Kirsten J. Sowers, Younghoon Park, Thavaleak Prum, Linette Rodriguez, Maria Ssozi, Jon Torres, Agnes A. Walsh, John E. Warner, Stephanie R. Weldon, Liling Xu, Kevin Wiehe, Max Crispin, Andrew B. Ward, Usha Nair, Beatrice H. Hahn, Dennis R. Burton, Lawrence Shapiro, Peter D. Kwong, Darrell J. Irvine, Raiees Andrabi, George M. Shaw, Facundo D. Batista

## Abstract

Current vaccine strategies to elicit broadly neutralizing antibodies (bnAbs) against HIV-1 generally propose complex, multi-boost immunization regimens. In rhesus macaques, SHIV infection has been observed to rapidly drive the development of some classes of bnAbs that share structural similarities with those in humans. Here, we generated a knockin mouse model with B cells bearing the unmutated common ancestor (UCA) of the V2 apex-targeted bnAb lineage, V033-a. A single immunization of mice with a germline-targeting native-like trimer was sufficient to recapitulate the ontogeny of the mature rhesus bnAb in knockin mice—including rare, disfavored somatic mutations—leading to the induction of antibodies that exhibited potent neutralization against both autologous and heterologous tier 2 viruses. A boost with Env escape mutant trimers further improved breadth and potency, and cryo-EM structure revealed the structural basis for heterologous neutralization breadth. Non-human primate and mouse models can thus combine with structure to serve as a platform for identifying and confirming immunogens that streamline HIV-vaccination regimens.

## Introduction

The remarkable diversity of circulating HIV-1 strains has proven a major challenge for vaccine development (Hemelaar, 2012). Nonetheless, the discovery of broadly neutralizing antibodies (bnAbs) in a subset of people living with HIV in recent decades has raised new hope for designing an effective HIV-1 vaccine (Burton et al., 2012; Burton & Hangartner, 2016; Kwong & Mascola, 2018). bnAbs arise following an initial prime of rare naive B cells with appropriate V-D-J features followed by a complex coevolutionary process between the escaping HIV-1 Env and antibody lineages undergoing affinity maturation (Doria-Rose & Landais, 2019; Liao et al., 2013; Rantalainen et al., 2018; Xiao et al., 2009). The most common strategy to reproduce this trajectory by vaccination, therefore, is similarly complex: immunization with a sequence of Env immunogens to selectively activate naïve B cells with specific features critical to bnAb epitope recognition, followed by further boosting to shepherd the response to acquire neutralization breadth (B. Briney et al., 2016; Dosenovic et al., 2015; Escolano et al., 2016; Jardine et al., 2013, 2015; Steichen et al., 2016). Priming immunogens designed for this approach are increasingly entering clinical trials, and some show promise (NCT06033209, NCT04224701, NCT05001373, NCT05471076) (Caniels et al., 2025; Willis et al., 2025)—the eOD-GT8 60mer, for example, induced precursors to VRC01-class bnAbs in 97% of vaccine recipients (Leggat et al., 2022). However, each step required for a complete vaccine series increases both the logistical challenges in delivery and the likelihood of incomplete immunization.

The expectation of extensive boosting regimens arises from the complex developmental pathways of bnAbs to many canonical epitope specificities, which often involves substantial acquisition of somatic hypermutation (SHM). VRC01-class bnAbs, which are specific to the CD4-binding site (CD4bs), are characterized by extreme SHM, with nucleotide variation from germline in the heavy chain of 20–35% and 15–20% in the light chain leading to amino acids identities of less than 50% (Zhou et al., 2013). In contrast, V2 apex bnAbs, which are commonly observed in HIV-infected human cohorts (Landais et al., 2016; Rusert et al., 2016; Walker et al., 2010) and in SHIV-infected macaques (Habib et al., co-submission), may require relatively low rates of SHM, may develop rapidly after priming of their germline unmutated common ancestors (UCA), and may be shepherded by only a few Env escape mutations (Bhiman et al., 2015; Doria-Rose et al., 2014; Habib et al., 2025; Landais et al., 2017). Therefore, it is thought that V2-apex bnAb UCAs may have simpler maturation pathways than bnAbs to other specificities.

Env Q23.17 (Poss & Overbaugh, 1999) has been identified in several screens of large virus panels as an Env sensitive to neutralization by germline-reverted V2-apex bnAbs, suggestive of an intrinsic germline-targeting property (Bonsignori et al., 2011; Gorman et al., 2016; Voss et al., 2017). This property is confirmed in an accompanying manuscript where we demonstrate that SHIV-Q23.17 is able to rapidly elicit V2-apex bnAbs in as many as 50% of infected macaques (Habib et al., 2025). While longitudinal lineage tracing of V2-apex directed antibodies from HIV-infected patients has been performed previously (Doria-Rose et al., 2014; Landais et al., 2017), lineage tracing in rhesus macaques provides greater opportunity to finely track bnAb development after infection via frequent sampling of B cells during bnAb maturation. One V2-apex lineage arising after SHIV-Q23.17 infection, V033-a, was chosen for further study because of its well-described ontogeny, marked neutralization breadth, potency, and the short timeframe in which it arose (Habib et al., 2025; Roark et al., 2024).

Immunoglobulin knock-in mouse models have been commonly deployed to examine candidate GT vaccine designs (Dosenovic et al., 2015; Huang et al., 2020; Jardine et al., 2015; Ray et al., 2024). However, even in favorable knockin mouse models, such vaccine strategies have required complex boosting regimens (B. Briney et al., 2016; Chen et al., 2021; Cottrell et al., 2024; Escolano et al., 2016; Haynes et al., 2023; Steichen et al., 2016; Tian et al., 2016; Wang et al., 2024; Xie et al., 2024). Therefore, we asked whether we could recapitulate the short ontogeny of the V033-a lineage in a knockin mouse model to expedite the elicitation of V2-apex bnAbs by vaccination. To do so, we knocked in an early precursor of the V033-a lineage (V033a-UCA I1) that differs from the phylogenetically inferred UCA by just one amino acid in the nontemplated junction between D and J (Habib et al., 2025)— generating what is, to the best of our knowledge, the first such macaquized immunoglobulin mouse model.

By using these models, we demonstrated that native-like stabilized HIV Env Q23.17 (Q23-APEX-GT1), can bind knockin V033a-UCA I1 B cells. We found that a single bolus priming immunization of Q23-APEX-GT1 was sufficient to select for critical mutations across the variable region of the B cell precursor lineage and endow isolated antibodies with notable neutralization breadth. Furthermore, a single boosting immunization resulted in additional gains in neutralization breadth and potency, with an escape-variant boost outperforming a homologous boost. Thus, we demonstrate that some near-native Envs are capable of priming and guiding the maturation of V2 apex bnAbs, that at least a subset of V2 apex bnAb UCAs are capable of rapidly acquiring substantial breadth and potency, and that the design of boosting immunogens based on Env escape may be a useful strategy to rapidly shepherd primed bnAb precursors into mature bnAbs, recapitulating the B cell evolution observed in the macaque after SHIV infection.

## Results

### Structure-guided design of native-like Q23-APEX-GT1 trimer for priming V2-apex bnAb B cell precursors

We previously identified a subset of primary HIV Envs that contain fewer potential N-linked glycosylation (PNG) sites in their V1V2 segments than do most other wildtype Envs, enabling V2 apex bnAb inferred germline B cell receptor (BCR) binding to native-like trimers (Andrabi et al., 2015; Gorman et al., 2016; Voss et al., 2017). These findings suggested that these native-like trimers could readily engage bnAb precursors at the V2 apex site without extensive protein engineering. More recently, in a rhesus macaque SHIV infection model, one such Env, Q23.17, consistently induced V2 apex bnAbs, underscoring its intrinsic ability to elicit site-targeted bnAb responses (Habib et al., 2025). Accordingly, in this study we sought to design a Q23.17 Env-based trimer immunogen that could be delivered across various platforms, including mRNA lipid nanoparticles (LNPs).

To generate prefusion-stabilized native-like Q23 Env trimers, we employed a strategy combining previously described NFL (native flexibly linked) and RnS (repair-and-stabilization) designs (Rutten et al., 2018; Sharma et al., 2015) with additional antibody-guided structure-based mutations. We optimized the trimer subdomains to enhance hydrophilic interactions at the apex, restrict CD4-induced conformational changes, and minimize off-target V3 epitope exposure. Stabilization targeted both gp120 (including V1V2 and V3 regions) and gp41 subunits (**Fig. 1A–B and Figs. S1&S2**). We systematically combined stabilized subdomain variants, covalently linked them with a glycine-serine linker, and assessed trimer formation using size-exclusion chromatography (SEC). The final construct, a covalently linked gp120-gp41 trimer, is referred to as a single-chain trimer (SCT). The initial design, Q23-SCT21, incorporated SOSIP mutations (501C-605C, I559P) and a disulfide-(DS) bond (201C-433C) to reduce CD4-induced epitope exposure (Sanders et al., 2013; Zhang et al., 2018). This core mutation set was retained across all ten variants (Q23-SCT21 – Q23-SCT30), with subsequent iterations introducing gp41 and gp120 stabilizing mutations (**Figs. S1&S2**).

**Figure 1.**
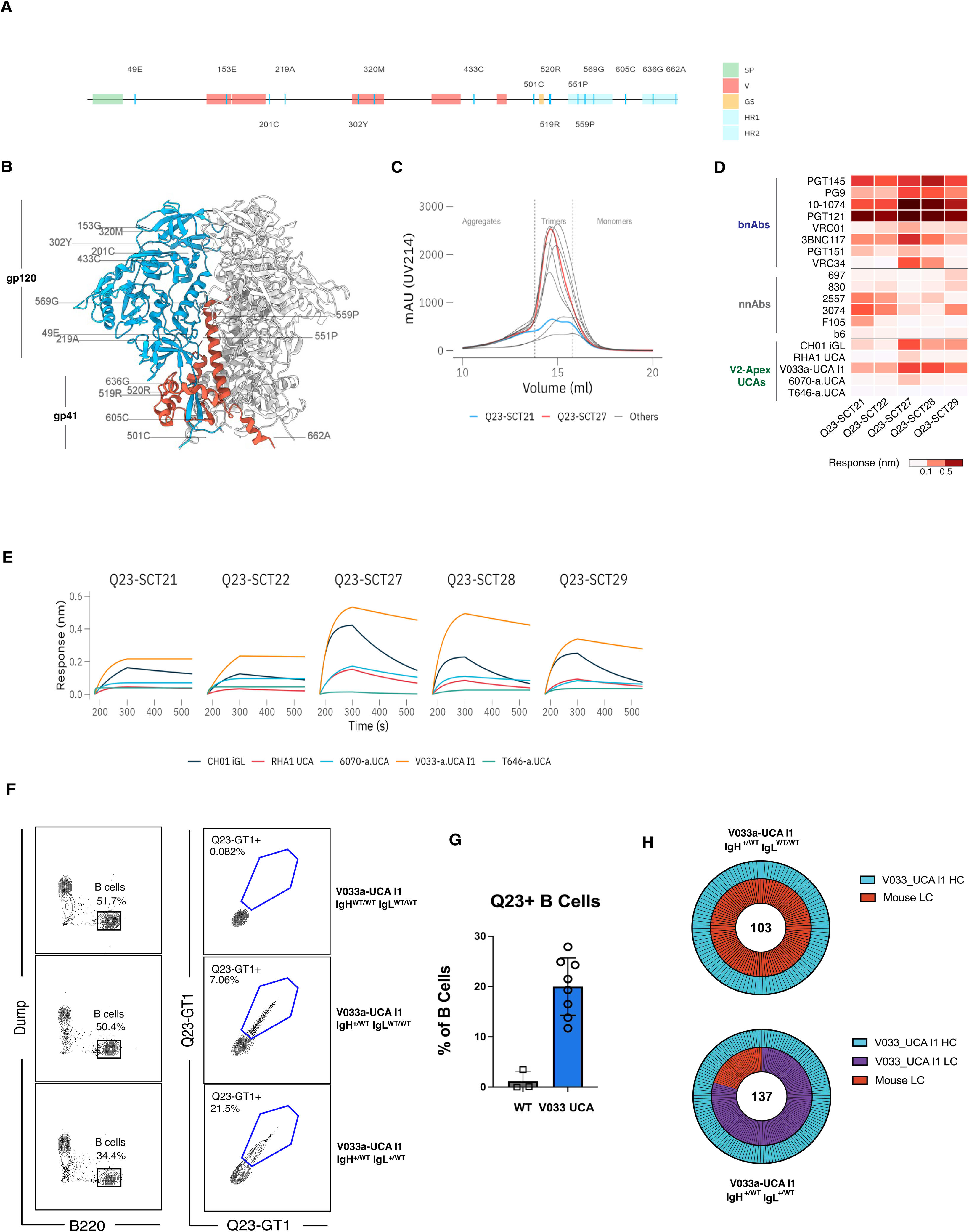
Generation and characterization of the Q23-SCT immunogen and V033a-UCA I1 mouse model. (A) Structure-guided mutations utilized in construct Q23-SCT27 are labelled and shown on a schematic map of HIV-1 Env. Signal peptide is shown in green, variable loops are indicated in red, flexible glycine-serine linker (G4S)2 is shown in yellow and HR1 and HR2 region within gp41 are shown in blue. (B) Amino acid substitutions in Q23-SCT27 are labelled and shown on the crystal structure of Q23 DS-SOSIP.664 trimer (PDB: 7LLK) with one protomer colored according to sub-regions (gp120 in blue and gp41 in red). (C) Representative size exclusion chromatography (SEC) profile of Galanthus-nivalis (GNL) purified Q23-SCTs. The fractions corresponding to aggregates, trimer and dimer/monomers are annotated. Fractions used for antigenic profiling are shown inside dotted lines. The lead candidate, Q23-SCT27, is shown in red while the base construct is shown in blue. All remaining constructs are shown in gray. (D) Antigenic profile of five SCTs against a small panel of bnAbs, unmutated common ancestors (UCA), inferred germline (iGL) and non-neutralizing antibodies was performed with bio-layer interferometry (BLI). Maximum response values were used for plotting heat map. (E) Kinetic curves from BLI for the five SCTs from panel D against the five RM V2 apex UCAs. V033 I1 showed highest stable binding against Q23-SCT27 to Q23-SCT29. (F) Representative FACS plots of peripheral B220^+^ B cell binding of Q23-SCT27 in C57BL6/J, V033a-UCA I1 IgH^+/WT^ IgL^WT/WT^ and V033a-UCA I1 IgH^+/WT^ IgL^+/WT^ mice. (G) Quantification of peripheral B cell binding of Q23-SCT27 in C57BL6/J and V033a-UCA I1 IgH^+/WT^ IgL^+/WT^. (H) Paired sequences of single-cell sorted Q23-APEX-GT1-binding naïve B cells V033a-UCA I1 IgH^+/WT^ IgL^WT/WT^ (upper) and V033a-UCA I1 IgH^+/WT^ IgL^+/WT^ (lower) mice (n=2 each). Outer circle shows heavy chain (HC) identity; inner indicates light chain (LC). V033a-UCA I1 HC = teal; V033A-UCA I1 LC = purple; native murine LC = red.

For gp41 stabilization, the Q23-SCT22 construct incorporated glycine substitutions at positions 569 and 636 to reduce helical flexibility (Guenaga et al., 2017), Q551P to rigidify gp41, and 519R-520R to decrease the fusion peptide’s hydrophobicity. The Q23-SCT23 construct included seven RnS-derived mutations (535N, 556P, 588E, 589V, 651F, 655I, 658V) (Rutten et al., 2018), while Q23-SCT24 combined all twelve gp41-stabilizing mutations from both Q23-SCT22 and Q23-SCT23. Lastly, Q23-SCT25 maintained these mutations but replaced the 501C-605C disulfide bond with a 501C-663C linkage (Yang et al., 2018). For gp120 stabilization, six additional mutations were introduced: 49E, 153E, 219A, 302Y, 320M, and 334S. The mutations 49E, 302Y, and 320M were derived from NFL-TD8 (Guenaga et al., 2015), 153E was selected based on structural analysis (PDB 7LLK) to stabilize the V2 apex, and 219A and 334S were adapted from RnS using the ADROITrimer (Rawi et al., 2020). These gp120 mutations were systematically incorporated into the following constructs: Q23-SCT26 = Q23-SCT21 + gp120 mutations, Q23-SCT27 = Q23-SCT22 + gp120 mutations, Q23-SCT28 = Q23-SCT23 + gp120 mutations, Q23-SCT29 = Q23-SCT24 + gp120 mutations, and Q23-SCT30 = Q23-SCT25 + gp120 mutations. This iterative design approach enhanced trimer stability and antigenic properties (**Figs. S1 & S2**).

SEC profiles of *Galanthus nivalis* lectin (GNL)-purified Q23-SCT soluble proteins revealed distinct peaks for aggregates, trimers, and dimer/monomers in most designs (**Fig. 1C**). Four Q23-SCT variants—Q23-SCT22, Q23-SCT27, Q23-SCT28, and Q23-SCT29—were selected based on higher trimer yield, sharper trimer peaks, and reduced aggregate or monomer/dimer peaks, indicating improved trimer assembly compared to the base construct (Q23-SCT21), which contained minimal mutations (**Fig. 1C and Fig. S1A&B**). The antigenic profile of the SCTs was evaluated using bio-layer interferometry (BLI) against a panel of HIV Env bnAbs, non-neutralizing antibodies (non-nAbs), and UCAs and iGLs of V2 apex bnAbs (**Fig. 1D**). Q23-SCT27 demonstrated enhanced BLI binding to several bnAbs and substantially reduced binding to V2i (697-30D and 830a) and CD4bs non-nAbs (b6 and F105), suggesting improved antigenic properties (**Fig. 1D**). Notably, Q23-SCT27 exhibited enhanced binding to several V2-apex bnAb UCAs and iGLs, including both human (CH01 iGL and PCT64 LMCA) and rhesus (RHA1 UCA and V033a-UCA I1). A comprehensive antigenic analysis of Q23-SCT27 with a larger panel of 70 monoclonal antibodies (mAbs), divided into bnAbs and non-nAbs, further confirmed its broad reactivity across different specificities of bnAbs (**Fig. S1C**). This binding was consistent with the mammalian cell surface-expressed trimer version of Q23-SCT27 (with wildtype Q23.17 transmembrane domain) and neutralization of the wild-type Q23.17 virus (**Fig. S3**). Of note, the cell surface-expressed trimer showed enhanced binding, likely due to avidity effects.

Negative stain electron microscopy (ns-EM) of Q23-SCT27 trimers revealed well-formed trimeric structures with distinct 2D-averaged classes, confirming the structural integrity of the engineered trimer (**Fig. S1D**). Proteomics-based site-specific glycan analysis (SSGA) of Q23-SCT27 showed a diverse glycan profile, including high mannose (green), complex glycans (pink), and unoccupied sites (gray) (**Fig. S1E**). Notably, no glycan signal could be resolved at PNGS 141 and 401, suggesting inherent flexibility not amenable to protease digestion. The well-formed trimeric structure and native-like glycan profile of Q23-SCT27, notably the conservation of oligomannose-type glycans at N160, validated its suitability as a vaccine candidate.

Based on these properties, we selected Q23-SCT27, henceforth called Q23-APEX-GT1 due to its enhanced binding to V2-apex bnAb UCAs and iGLs, as our lead candidate for preclinical testing.

### Generation of knockin mice expressing a precursor to a rhesus macaque bnAb

To model Q23-APEX-GT1 engagement in a stringent pre-clinical mouse model, we used our established CRISPR/Cas9-mediated knockin method (Lin et al., 2018; Wang et al., 2021) to introduce the heavy and light chains of the V033-a lineage UCA designated V033a-UCA I1. The naïve germline precursor, or UCA, was derived from analyses of early sequential IgM+ naïve B cell and IgG+ memory B cell immunoglobulin sequences from the peripheral blood of macaque V033. Phylogenetic analysis of these sequences coalesced to two heavy chains that differed by a single amino acid in the nontemplated D-J junction (Habib et al., 2025). There was also a single amino acid difference in VL. One of paired VH and VL sequences, we designated V033-a.UCA; the other, we designated V033a-UCA I1. For construction of the KI mouse lines, we used V033a-UCA I1 as it exhibited a higher binding affinity and greater neutralization potency against the Q23.17 Env; furthermore, we could determine its high resolution cryoEM structure in complex with the Q23.17 Env trimer (Habib et al., 2025). The successful knockin was confirmed through genotyping and is referred to as the V033a-UCA I1 mouse below. Peripheral B cells from V033a-UCA I1 mice bound Q23-APEX-GT1 trimer conjugated to streptavidin probes in the presence or absence of V033A-UCA I1 knockin light chain (**Fig. 1F**); for V033a-UCA I1 IgH^+/WT^ IgL^+/WT^, ∼20% of peripheral B cells bound the probe (**Fig. 1G**). BCR sequencing of Q23-APEX-GT1-binding B cells using 10x Genomics revealed that V033a-UCA I1 heavy chain (HC) paired with multiple murine light chains (LC) in V033a-UCA I1 IgH^+/WT^ IgL^WT/WT^ mice, whereas in V033a-UCA I1 IgH^+/WT^ IgL^+/WT^ mice, the majority of knockin HCs paired with knockin LCs (**Fig. 1H**). These results confirmed the successful knockin of macaque V033a-UCA I1 into a murine model: V033a-UCA I1 B cells were developmentally normal and bound the Q23-APEX-GT1 trimer.

### Q23-APEX-GT1 protein trimer and Q23-APEX-GT1 saRNA efficiently activate VO33a-UCA I1 B cells

To determine whether the Q23-APEX-GT1 trimer could engage and activate V033a-UCA I1 B cells in vivo, we transferred V033a-UCA I1 IgH^+/WT^ IgL^+/WT^ CD45.2^+/+^ B cells intravenously (IV) into recipient CD45.1^+/+^ C57BL/6J to establish a frequency of 8 CD45.2s in 10^6^ total B cells (Lin et al., 2018; Melzi et al., 2022). One day later, adoptively transferred animals were immunized subcutaneously (SC) in the base of the tail with 10 µg of Q23-APEX-GT1 prepared in 5 µg of saponin/MPLA nanoparticles (SMNP) adjuvant (Silva et al., 2021) (**Fig. S4A**). The immune responses in the draining inguinal lymph nodes were analyzed at weeks 2 and 4 post-immunization (wpi). Immunized animals at 2 wpi were able to form germinal centers (GCs) (Dump^-^ B220+ CD38^lo^ CD95^+^ among live cells) and these GCs were sustained up to 4 wpi (**Fig. S4B&C**). Notably, a substantial fraction of these GCs was composed of adoptively transferred CD45.2 V033a-UCA I1 B cells, with ∼40% binding Q23-APEX-GT1 (**Fig. S4B&D**), demonstrating successful activation of knockin B cells and GC recruitment by Q23-APEX-GT1 protein trimer priming.

The low precursor frequency of V2-apex bnAb UCAs is one of the principal barriers to their elicitation (Mishra et. al, *Immunity* in review) (B. S. Briney et al., 2012; Habib et al., 2025; Willis et al., 2022). To recapitulate this in a model of immunization, we increased the stringency of the model by transferring V033a-UCA I1 IgH^+/WT^ IgL^+/WT^ CD45.2^+/+^ B cells IV into recipient CD45.1^+/+^ C57BL/6J mice (Lin et al., 2018; Melzi et al., 2022). One day later, animals were immunized with 10 µg of Q23-APEX-GT1 prepared in 5 µg of SMNP adjuvant subcutaneously (**Fig. 2A**). Analysis of draining lymph nodes in immunized animals showed strong early GC responses (Dump^-^ B220+ CD38^lo^ CD95^+^ among live cells) with Q23-APEX-GT1 trimer, which diminished over time (**Fig. 2B&D**). V033a-UCA I1 CD45.2 B cells resided in GCs at a high percentage up to 6 wpi with ∼40% in week 2, ∼60% week 4 and ∼40% in week 6 (**Fig. 2B&D**). Immunization with Q23-APEX-GT1 trimer, therefore, elicited long-lasting precursor participation in GCs despite stringent initial frequencies.

**Figure 2.**
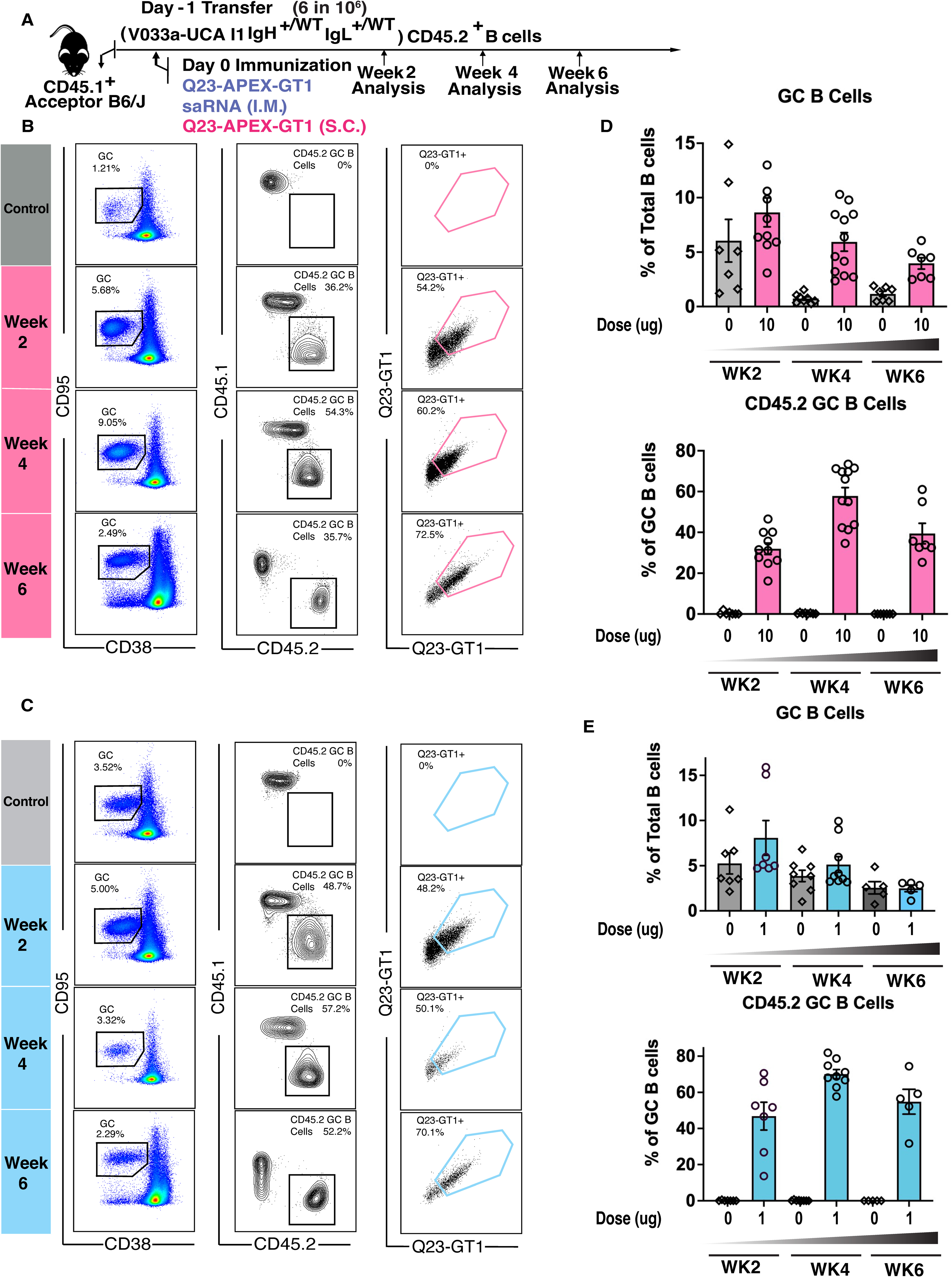
Q23-APEX-GT1 can prime V033a-UCA I1 B cells at low frequencies. (A) Schematic of mouse adoptive transfer and immunization experiments. WT CD45.1^+/+^ mice received V033a-UCA I1 CD45.2^+/+^ B cells through intravenous transfer one day prior to immunization with 1 μg per hindlimb (total 2 μg) of saRNA-LNP delivered intramuscularly (IM) or 10 μg of Q23-APEX-GT1 trimer adjuvanted with saponin/MPLA nanoparticles (SMNP) subcutaneously (SC). SMNP adjuvant without trimer was used as the control for protein immunizations and LNPs containing an unrelated saRNA for the Q23-APEX-GT1 saRNA LNP immunizations (B, C) Representative FACS plots showing GCs, CD45.2 B cells in GCs, and their binding to Q23-APEX-GT1 during weeks 2, 4 and 6 post-immunization with Q23-APEX-GT1 trimer with SMNP (B) or LNPs containing Q23-APEX-GT1 saRNA (C). (D) Quantification of GC B cells (upper) and CD45.2 V033a-UCA I1 B cells in GCs (lower) in post-immunization by Q23-APEX-GT1 adjuvanted with SMNP (denoted as 10) or SMNP alone (0). Each group sums three independent immunizations. (total n=7–10). (E) Quantification of GC B cells (upper) and CD45.2 V033a-UCA I1 B cells in GCs (lower) in responses post Q23-APEX-GT1 saRNA LNP (denoted 1) or empty LNP immunization (0). Each group sums three independent immunizations (total n=7–10).

For comparison with protein-based immunization, we developed a self-amplifying RNA (saRNA) (Melo et al., 2019), encoding a membrane-anchored form of Q23-APEX-GT1 utilizing the transmembrane domain of wildtype Q23.17 virus. To determine whether membrane-anchored Q23-APEX-GT1 delivered by saRNA could activate V033 precursors, we performed adoptive transfer as described above and immunized the recipients with Q23-APEX-GT1 saRNA containing LNPs intramuscularly (IM) **(Fig. 2A).** Activation and GC participation was similar to that obtained by trimer immunization out to week 4 (**Fig. 2C&E)**. Thus, Q23-APEX-GT1, delivered as either a trimer protein with SMNP adjuvant or saRNA LNP encoding membrane-bound trimer, effectively activates V033a-UCA I1 B cells and supports Q23-APEX-GT1-specific durable germinal center responses.

### A single Q23-APEX-GT1 immunization leads to substantial on-track V033-a mutations

We next sought to understand the degree to which a single immunization was able to recapitulate V033-a lineage ontogeny as observed in the macaque. As we observed a substantial number of antigen-positive CD45.2^+^ B cells participating in GCs after immunization, we isolated Q23-APEX-GT1^+^ CD45.2 B cells after protein-trimer immunization (**Fig. 2B&C**) to determine the extent and quality of clonal expansion and SHM. Analysis of V033a-UCA I1 knockin BCRs in paired sequencing data revealed substantial diversification in the heavy chains (**Fig. 3A**). Furthermore, several amino acid replacements that were observed in the rhesus macaque V033 over the course of the maturation of the V033-a lineage appeared as early as 2 weeks post-immunization (**Fig. 3A**), and on-track mutations accumulated over time (**Fig. 3B&C**) (Habib et al., 2025). Furthermore, the heavy chain CDR3, which is important for V2-apex recognition, displayed rapid clonal divergence over time (**Fig. 3A–C, middle panel**). During the course of immunization, the knockin heavy chain of the adoptively transferred B cells displayed a preferential association with the knockin light chain (**Fig. 3A–C**); both heavy and light chains accumulated mutations rapidly over time after just a single bolus priming immunization (**Fig. 3D**). As with heavy chains, knockin light chains displayed “on-track” V033 mutations (**Fig. 3E**). Similar to protein trimer immunization, BCRs sequenced after saRNA immunization also demonstrated a rapid gain of SHM in V033a-UCA I1 HCs **(Fig. S6A).**

**Figure 3.**
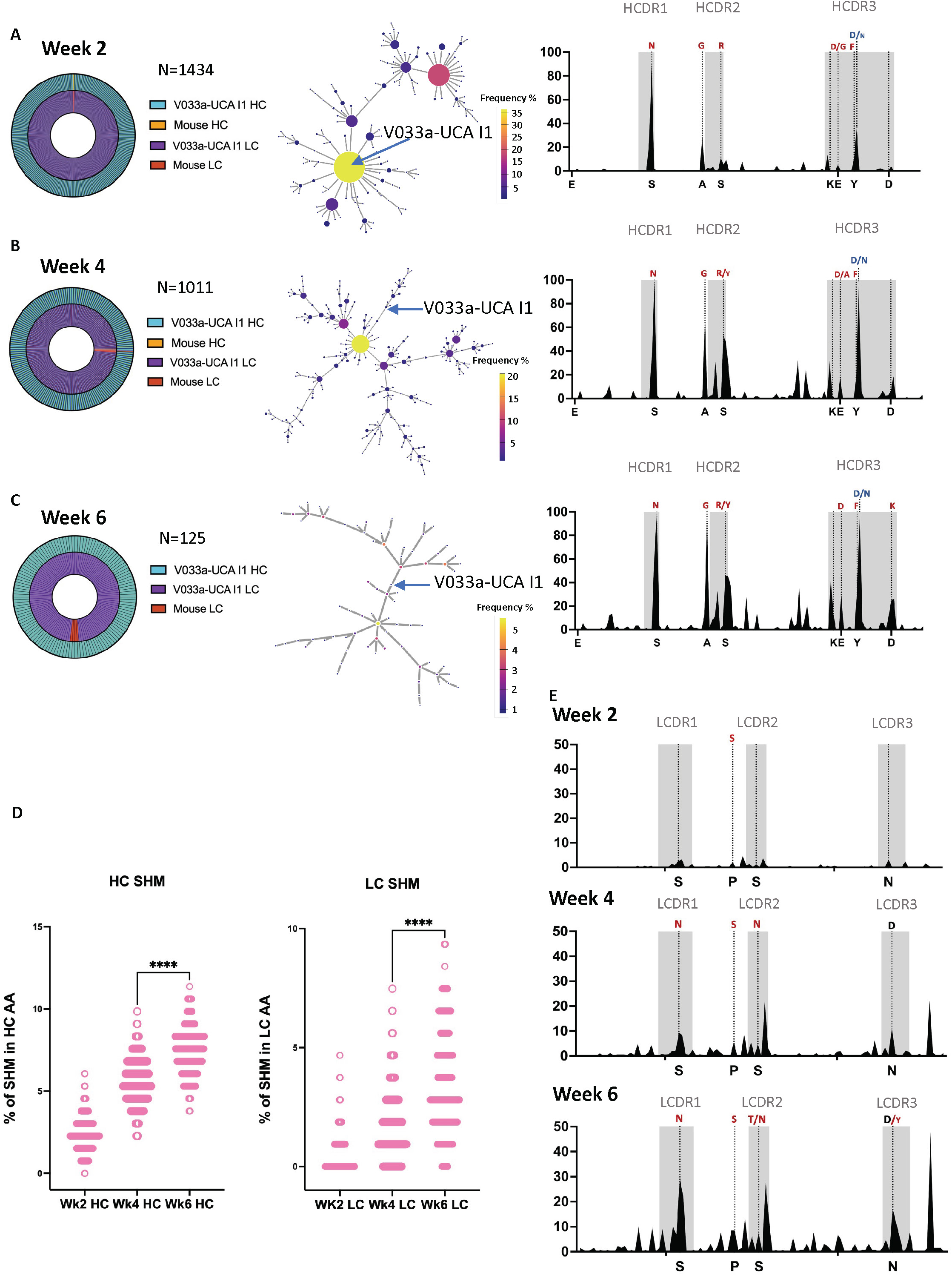
Priming leads to recapitulation of V033 ontogeny and rapid gain of neutralization breadth. (A, B, C) (Left) HC (outer) and LC (inner) usage by single cells from Q23-APEX-GT1-specific CD45.2^+^ sorted 2 (A), 4 (B), and 6 (C) weeks post-immunization with Q23-APEX-GT1 trimer adjuvanted with SMNP. (Center) Divergence of knockin V033a-UCA I1 heavy chain post immunization represented through phylogenetic trees. (Right) HC mutation frequencies; selected mutations in V033a-UCA I1 HC present in mature V033 lineage bnAbs are represented in red and mutations found at intermediate stages of V033 lineage development (Habib et al., 2025) are marked in blue. (D) Total amino acid (AA) mutations in V033a-UCA I1 IGHV and IGKV at weeks 2, 4, 6 post immunization. (E) LC mutation frequencies after 2, 4 and 6 weeks after immunization with Q23-APEX-GT1 trimer. Select mutations in present in mature V033 lineage bnAbs are represented in red; mutations found at intermediate stage of V033 lineage development is marked in blue.

The rapid clonal divergence observed and the accumulation of “on track” mutations led us to further characterize a subset of the clones from 4 wpi and 6 wpi to express as monoclonal antibodies (mAbs). All of the 6 wpi prime-derived mAbs had high affinity for the immunogen Q23-APEX-GT1 (**Fig. S7A**). We therefore tested the efficacy of these antibodies against a limited panel of HIV-1 strains in a pseudovirus neutralization assay. All of the selected mAbs could neutralize the autologous Q23.17 virus with IC_50_ values ranging from 0.029–5.2 µg/ml. Some mAbs could neutralize heterologous viruses that are sensitive to V2-apex bnAb neutralization (**Fig. S7B**), albeit with a lesser potency than the mature V033-a.01 mAb. This subset of viruses was the first one to become sensitive to neutralization from V033 plasma in the NHP (Habib et al., 2025). Three of the selected mAbs from 4 wpi neutralized up to 43% of strains in a larger panel of heterologous tier 2 viruses (**Fig. S7C**). A single Q23-APEX-GT1 immunization can thus lead to the acquisition of substantial neutralization breadth.

### Neutralization breadth and potency attained late in single immunization-derived antibodies

Env-targeting bnAbs may be enriched for mutations that are less favored due to codon divergence and the ability to mutate when the residue is in AID “cold spots” (Wiehe et al., 2018). As we observed that “on-track” mutations accumulated over time in V033a-UCA I1 knockin BCRs, particularly in the CDRs, we sought to determine whether an increase in less favored mutations over the course of a longer GC residency could enhance potency and breadth. ARMADiLLO analysis of HCs isolated from rhesus macaques post-SHIV infection predicted that rare mutations appear in the mature V033 lineage with greater frequency over time **(Fig. S5)**. We used ARMADiLLO to select antibodies from 6 wpi enriched for less favored mutations, some of which appear only in week 6 after priming **(Fig. S6B)**. The mAbs derived from 6 wpi were able to neutralize the autologous Q23.17 virus, as well as the first Env escape variant detected in the macaque, N187S, with higher potency (**Fig. 4A**) and demonstrated heterologous neutralization capacity in a limited panel (**Fig. 4A**). In a larger panel of heterologous tier 2 virus strains, one of these mAbs, T6-P_H03, showed neutralization breadth approaching that of the mature V033 (**Fig. 4B**). Additionally, all of the 6 wpi-derived antibodies bound with high affinity to the escape variants of Q23 and other V2-Apex bnAb sensitive Envs (**Fig. 4C**). We furthermore observed that, after priming with either protein trimer or saRNA, Q23-APEX-GT1-N187S binding can be observed inside the GCs as early as 4 wpi and continues to mature through 6 wpi **(Fig. S8B&C).** However, we observed only weak serum neutralization at later time points in the subset screened and tested after priming (4 wpi and 6 wpi), and only in animals that received Q23-APEX-GT1 trimer adjuvanted with SMNP **(Fig. S9A&B).** Serum neutralization, therefore, was not extensive after a single immunization, but mAb affinity, neutralization, and B cell binding data are nonetheless consistent with the development of a response to the escape variant (N187S) early after bolus priming.

**Figure 4.**
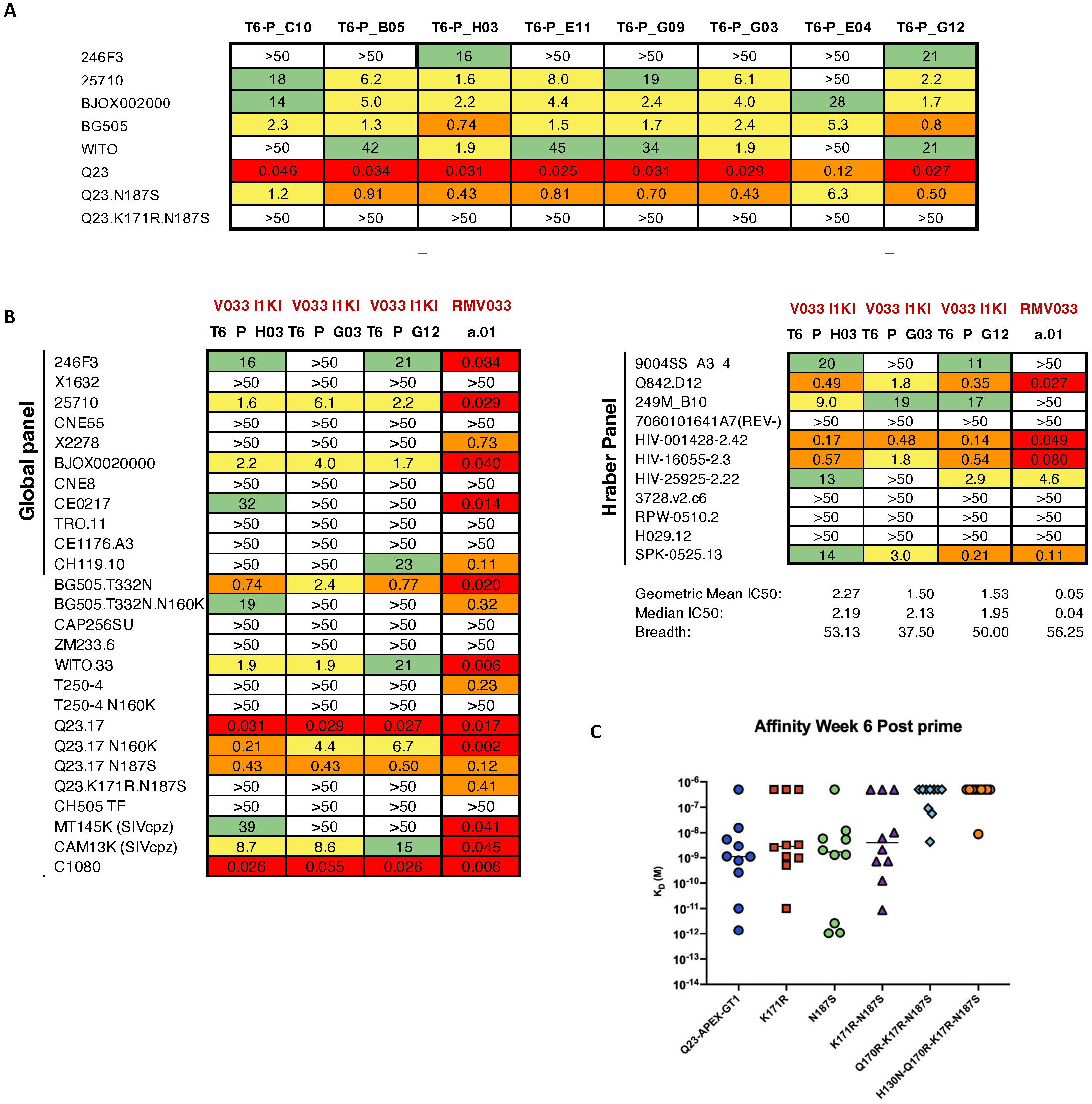
Neutralization breadth and potency of antibodies derived post-prime. (A) IC50 values of antibodies from week 6 post-priming by the protein trimer against autologous (Q23), escape variants (Q23.N187S, Q23.K171R.N187S) and limited panel of 5 heterologous tier-2 HIV-1 strains. Nomenclature: T = time in weeks; P = Prime; final alphanumeric triplet = mAb identity. (B) IC50 values of three post-prime antibodies from (A) against a 37-member panel of HIV-1 strains, including 11 from the Tier-2 Global panel and 11 from the Hraber panel; macaque mature V033 (RM V033 P2A6) shown for comparison (C) BLI affinity values of antibodies derived week 6 post-protein-prime against autologous Q23-APEX-GT1 and escape-variant Envs.

To investigate the molecular basis for heterologous neutralization breadth elicited by the single priming immunization, we determined the structure of the T6_P_H03 antigen-binding fragment (Fab) in complex with Q23-APEX-GT1 using single particle cryogenic electron microscopy (cryo-EM) (**Fig. 5, Figs. S10&S13, Table S1**). We previously found that, unlike many V2 apex bnAbs, the V033-a lineage can bind 3 Fabs per envelope trimer owing to (i) the ability to recognize monomeric gp120 and (ii) an HCDR3 which does not extend past the C-strand and into the middle of the envelope trimer, which would otherwise clash in the context of multiple Fabs (Roark et al., 2024)(Mishra et al., *Immunity* in review). Here, we obtained a 3D cryo-EM reconstruction of the T6_P_H03 complex that extended to 3.5 Å resolution and similarly revealed three Fabs bound to a single trimer, with an angle of approach and orientation nearly identical to V033a-UCA I1 (**Fig. 5A, Fig. S10**) (Habib et al., 2025). The elongated T6_P_H03 HCDR3 beta-strand precisely superimposed with that of V033a-UCA I1 and replicated the V033-a class-defining antiparallel strand-strand interaction with the V2 apex C-strand, resulting in the same three mainchain hydrogen bonds with the amide and carbonyl of R169 and the amide of K171. T6_P_H03 almost exclusively acquired heavy chain SHM in response to Q23-APEX-GT1 immunization (12 heavy chain residues vs. 1 light chain residue) (**Fig. 5B**), which is consistent with the heavy chain-dominated mode of recognition exhibited by the V033-a lineage (Habib et al., 2025; Roark et al., 2024). T6_P_H03 had acquired SHM at 8 of 12 identical positions observed in the latest V033-a lineage developmental intermediate (V033-a.I6), half of which fell within the Fab interactive surface (**Fig. 5C**). SHM in 5 of 8 conserved positions yielded residues identical to one or more developmental intermediates (V033-a.I2-6), including 3 paratope residues that engaged the C-strand more favorably (**Fig. 5D**). V033a-UCA I1 utilizes two consecutive Tyr residues, Y100_C_ and Y100_D_, to stabilize the extended aliphatic chains of C-strand residues R169 and K168, respectively. In contrast, T6_P_H03 acquired: F100_C_, which may be more sterically favorable since the hydroxyl group inserting between N156 and R169 is removed; and D100_D_, which forms a salt bridge with K168 (**Fig. 5D, left**). Further, both V033a-UCA I1 and T6_P_H03 engage the terminal amine group and extended aliphatic chain of C-strand residue K171 through D99 and Y100, respectively; however, T6_P_H03 acquired N31 through SHM, which forms an additional hydrogen-bond with K171 (**Fig. 5D, right)**.

**Figure 5.**
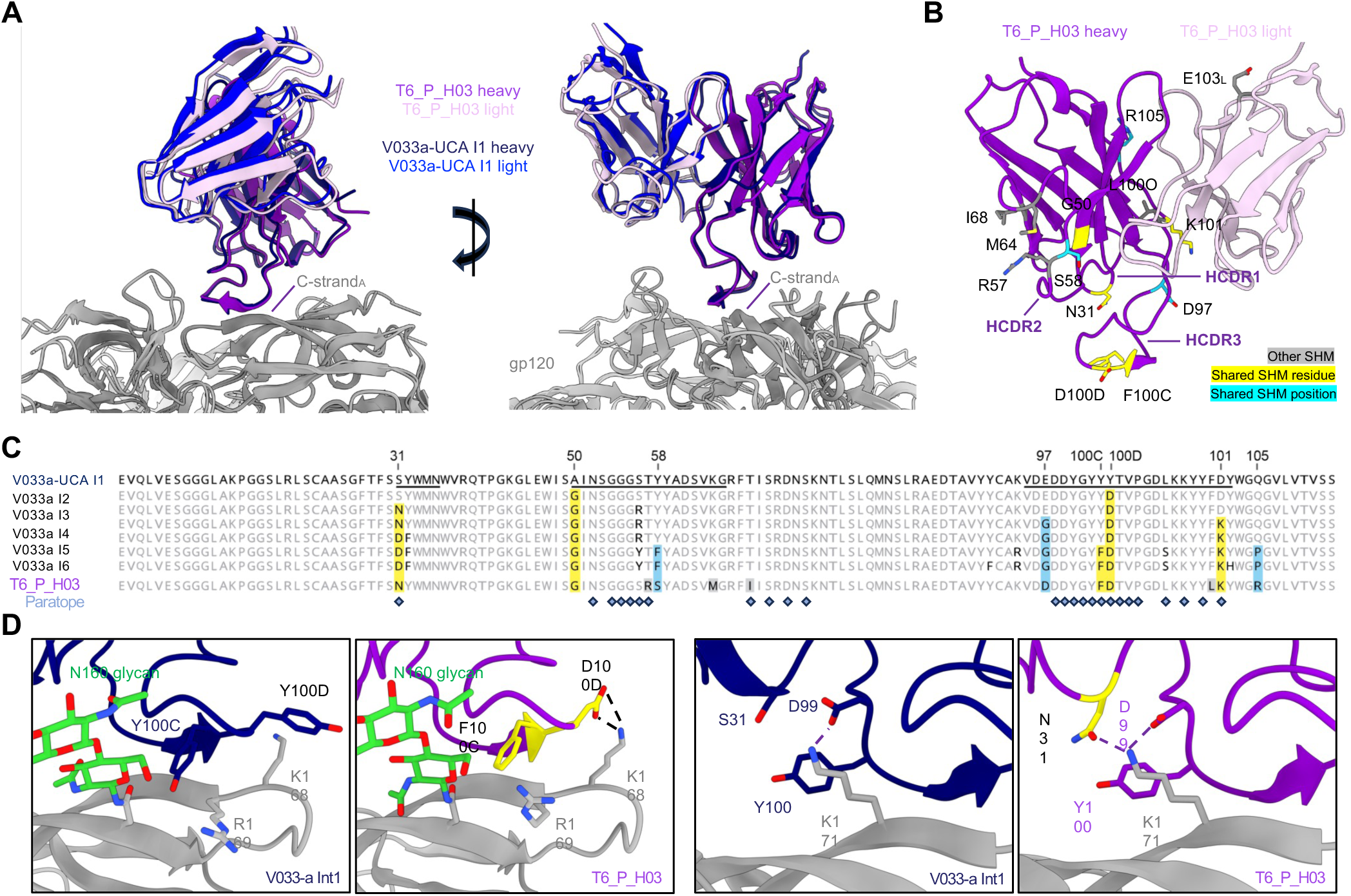
Structural basis of heterologous neutralization breadth for prime-derived antibody T6_P_H03. (A) Orthogonal views for the gp120 alignment of V033-a I1 and T6_P_H03 cryo-EM structures in complex with Env SOSIP trimer. Only one Fab per trimer is shown for clarity. (B) Visualization of SHM residues in the T6_P_H03 Fab structure. Mutated residues are shown in stick representation and colored according to legend: yellow if the mutation is identical to one or more V033-a developmental intermediates; teal if only the mutated position is shared, not the resulting residue, with the V033-a developmental intermediates; and gray if the mutation if unique to T6_P_H03. The label for the site of light chain mutation is denoted with an “L” subscript, (C) Top; Heavy chain residue alignment of V033-a lineage developmental intermediates (I1-6), the matured broadly neutralizing antibody V033-a.01, and T6_P_H03. The V033-a I1 sequence is designated as the reference, and matching unmutated germline residues in all other sequences are depicted in light gray. The HCDRs are underlined in the I1 sequence. Conserved patterns of somatic hypermutation between T6_P_H03 and one or more of the developmental intermediates are highlighted similarly to panel B. Paratope residues from the Prime_H03 structure are designated with slate gray diamonds. (D) Left; comparison of interactions for heavy chain paratope residues 100C and 100D in V033-a I1 and T6_P_H03. Right; comparison of C-strand K171 interactions for heavy chain epitope residues 31, 99, and 100 in V033-a I1 and T6_P_H03. Yellow residues are from panel B and C.

Thus, with a single immunization, we were able to induce antibodies capable of binding autologous Env escape variants and neutralizing heterologous tier 2 viruses. Together, these data reveal that a single immunization with Q23-APEX-GT1 in mice can recapitulate multiple immunogenetic and structural maturation pathways leading to the acquisition of breadth in the SHIV-elicited V033-a bnAb lineage.

### Homologous Q23-APEX-GT1 boosting increases serum neutralization breadth and on-track mutations

The ability of the native-like Q23-APEX-GT1 trimer to elicit a broadly reactive response in these mice raised the possibility that a homologous boost might drive these precursors to acquire the breadth and potency observed in the V033 macaque (Habib et al., 2025; Roark et al., 2024), and potentially serum neutralization, obviating the need for an extended immunization series. We therefore delivered a homologous boost to the recipient CD45.1^+/+^ C57BL/6J (donor V033a-UCA I1 IgH^+/WT^ IgL^+/WT^ CD45.2^+/+^ B cells) mice 65 days after the initial priming with Q23-APEX-GT1; at week 3 post-boost, the draining lymph nodes were analyzed for GC formation (**Fig. 6A**). The homologous boost could perpetuate the GCs and CD45.2 V033 B cells remained capable of binding both the priming immunogen as well as Q23-APEX-GT1 N187S, the next variant emerging from SHIV-Q23.17 escape in the V033-source (**Fig. 6B**) (Habib et al., 2025). We observed a several fold-increase in autologous Q23.17 serum neutralization and neutralization of an N160 glycan knockout variant after homologous boosting. Although we observed serum neutralization in one of the mice, Q23.17N187S was not sensitive to neutralization by sera from most animals (**Fig. 6C**). Nonetheless, serum neutralization was enhanced relative to prime-only.

**Figure 6.**
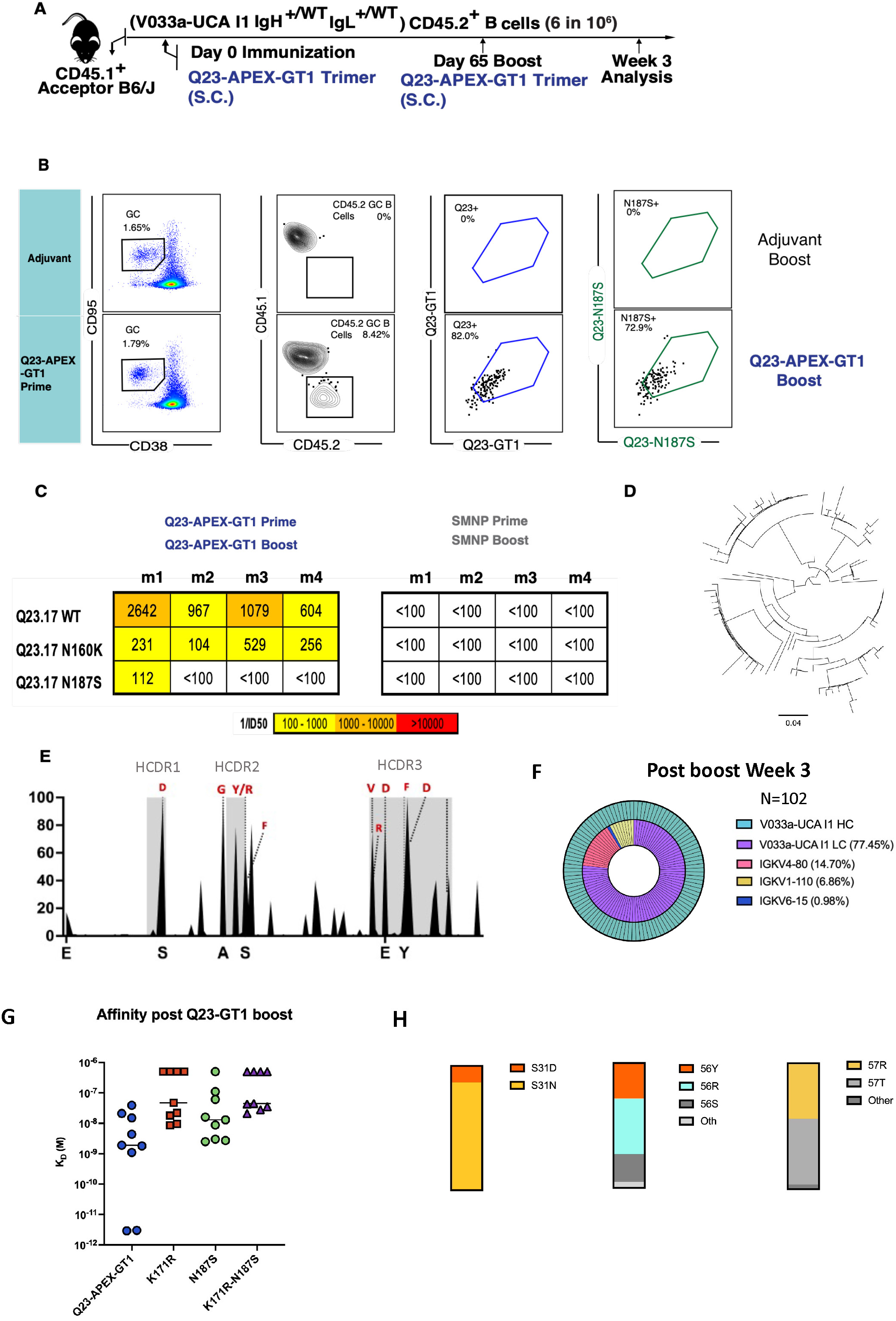
Homologous boosting leads to serum neutralization breadth and further on-track mutations. (A) Schematic presentation of mouse adoptive transfer and immunization experiments. WT CD45.1^+/+^ mice received V033a-UCA I1 B cells through intravenous transfer one day prior to immunization by Q23-APEX-GT1 trimer adjuvanted with SMNP. Day 65 post prime animals were boosted with Q23-APEX-GT1 adjuvanted with SMNP. Response was analyzed three weeks post immunization. (B) Representative FACS plots showing GC, CD45.2 B cells in GCs and their binding to Q23-APEX-GT1 and Q23-N187S during week three post-immunization with Q23-APEX-GT1 trimer with SMNP. Prime-boost with SMNP adjuvant without trimer was used as control for protein immunizations. (C) Reverse ID50 of serum of animals receiving Q23-APEX-GT1 Prime and Q23-APEX-GT1 boost week 3 post immunization against Q23.17, Q23.17 N160K and Q23.17 N187S escape variant. (D) Divergence of knockin V033a-UCA I1 heavy chain post boost represented through phylogenetic tree. (E) HC mutation frequencies at 3 weeks post-immunization by V033a-UCA I1 KI HC. Selected mutations in V033a-UCA I1 HC present in mature V033 lineage bnAbs are represented in red. (F) HC (outer) and LC (inner) usage by single cells from Q23-APEX-GT1-specific CD45.2^+^ B cells sorted three weeks post-boost. (G) BLI binding values and affinity of antibodies derived 6 weeks post-prime or 3 weeks post-boost against autologous Q23 Env and escape variant Envs. (H) Frequency of selected rare mutations in HCDR1 and HCDR2 post Q23-APEX-GT1 trimer prime and Q23-APEX-GT1 Boost.

We next sought to dissect the components of this serum response by examining SHM and affinity maturation in post-boost CD45.2^+^Ag^+^GC BCRs. As the prime-derived mAbs which could neutralize Q23.17 N187S with higher potency were overall better at neutralizing heterologous tier-2 viruses, we used Q23-APEX-GT1 N187S as a bait for sorting to enrich for potentially more potent mAbs. Sorted cells from homologous boosts showed further divergence of heavy chain (**Fig. 6D**), including a greater percentage of on-track mutations than previously observed after priming alone (**Fig. 6E**). Sorted cells also showed a higher preference for endogenous mouse light chains than before (**Fig. 6F**). Relative to prime-derived mAbs, there was similar affinity maturation even after receiving a boost (**Fig. 6G**). Three ARMADiLLO-predicted mutations were found to increase, as well (**Fig. 6H**). Thus, homologous boost-derived antibodies demonstrated greater serum neutralization breadth than animals that received bolus prime immunization and in one instance could accommodate Q23.17 escape variants detected in the macaque, despite having never encountered them *in vivo*.

### Escape variant boosting leads to higher SHM, including on-track mutations

In natural infections, bnAbs emerge from the coevolutionary process between viral escape and the host immune response (Doria-Rose & Landais, 2019; Liao et al., 2013; Rantalainen et al., 2018; Xiao et al., 2009); we therefore next asked whether Env escape mutations identified *in vivo* could be incorporated into boosting immunogens to drive bnAb maturation (Roark et al., 2021, 2024). In the post-prime germinal centers, the V033a-UCA I1 B cells bound to the escape variant Q23-APEX-N187S from 4 wpi (**Fig. S8**); we therefore immunized animals primed with Q23-APEX-GT1 with the escape variant Q23-APEX-GT1-N187S on day 65 (**Fig. 7A**). Serum neutralization of both autologous Q23.17 and escape variant N187S increased after Q23-APEX-GT1 N187S boost, whereas sensitivity to N160 glycan removal varied by animal (**Fig. S11**). Q23-APEX-GT1 N187S-boosted mice retained fewer CD45.2 B cells in the GC than those receiving a homologous Q23-APEX-GT1 boost (**Fig. 7B&C**). Cells were then sorted with Q23-APEX-GT1 N187S probe to select for antibodies capable of binding the escape variants, HCs recovered after the Q23-APEX-GT1 N187S boost were more mutated than those arising from the Q23-APEX-GT1 boost (**Fig. 7D**). Hotspot analysis also showed that a rare lysine mutation in the HCDR3, which was observed 6 wpi after prime, was enriched in the variant-boosted animals but not in those which received the homologous boost (**Fig. 7E**). BCRs from CD45.2^+^Ag^+^GC B cells isolated from mice immunized by the escape variant boost used endogenous mouse LCs at low rates **(Fig. S11B).** In LCs, the Q23-APEX-GT1-N187S boost did not increase overall mutation rates but did select for both increased frequency of previously observed mutations as well as additional lineage-recapitulating mutations **(Fig. S11C&D).** Thus, boosting with Q23-escape variants increased SHM overall, enriching on-track mutations.

**Figure 7.**
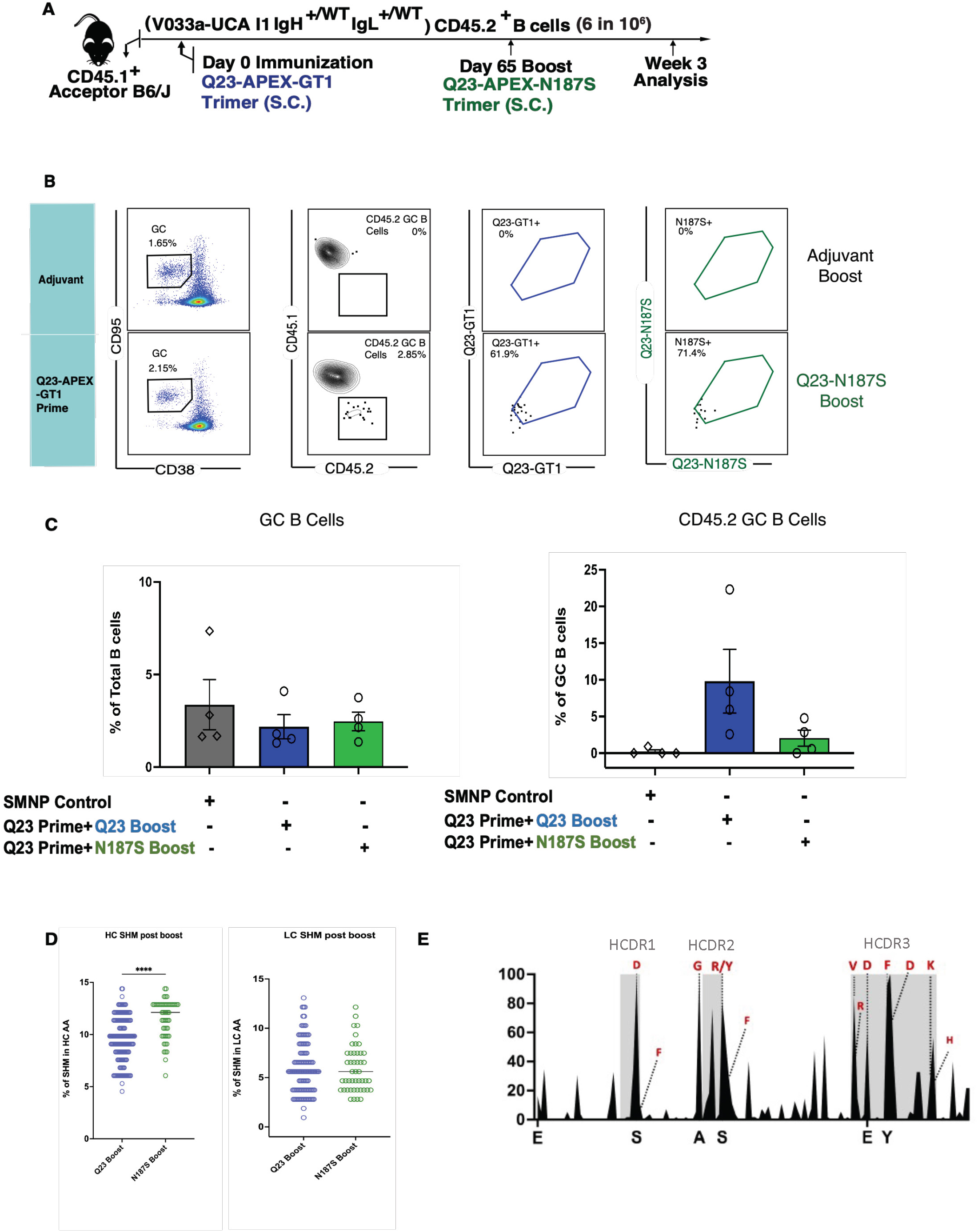
Escape-variant Env boosting leads to increased SHM and more on-track mutations. (A) Schematic presentation of mouse adoptive transfer and immunization experiments. Mice received V033a-UCA I1 B cells through intravenous transfer one day prior to immunization by Q23-APEX-GT1 trimer adjuvanted with SMNP. Prime-boost immunization by SMNP adjuvant without trimer was used as a control. Day 65 post-prime animals were boosted with Q23-N187S adjuvanted with SMNP. Response was analyzed 3 weeks post immunization. (B) Representative FACS plots showing GC, CD45.2 B cells in GCs and their binding to Q23-APEX-GT1 and Q23-N187S 3 weeks post-boost. (C) Quantification of GC B cells (left) and CD45.2 V033A-UCA I1 B cells in GCs (right) post-boost. (D) Total amino acid (AA) mutations in V033a-UCA I1 IGHV 3 weeks post-homologous (blue) or escape-variant (green) boost, compared with prime-only (gold) at weeks two, four, and six post-prime. (E) V033a-UCA I1 KI HC mutation frequencies week three post-boost. Select mutations present in mature V033-lineage bnAbs are represented in red.

### Both homologous and escape variant boosting enhance breadth and potency

To determine whether this overall increase in SHM was associated with a change in functionality, we expressed mAbs from animals receiving both homologous and variant boosts and subjected them to similar pseudovirus neutralization assays. mAbs were selected on the basis of either: 1) mutation frequency, 2) structure (CDRH3 stabilization), or the 3) presence of rare mutations. All boost-derived mAbs neutralized autologous Q23.17 virus and more potently neutralized Q23.17-N187S than mAbs derived post-prime (**Fig. 8A**). The neutralization ability of these antibodies was dependent to a varying degree on N160 glycan removal, but the antibodies were substantially more sensitive to C-strand mutations. Similarly, in a larger heterologous virus panel, Q23-APEX-GT1-boosted antibodies could neutralize WT Q23.17 and a large panel of heterologous viruses with a higher potency than prime-derived antibodies (**Fig. 8B**). The N187S variant boost-derived G05 antibody showed slightly greater neutralization breadth but also greater potency (**Fig. 8B**). The boost-derived antibodies also demonstrated enhanced binding to Q23 escape variants, with the G05 mAb derived from the N187S boost being the most potent in both neutralization and binding (**Fig. 8B&C**). Thus, some mAbs derived post-boost displayed improved neutralization and trimer binding and boosted animals displayed much higher serum neutralization than prime only.

**Figure 8.**
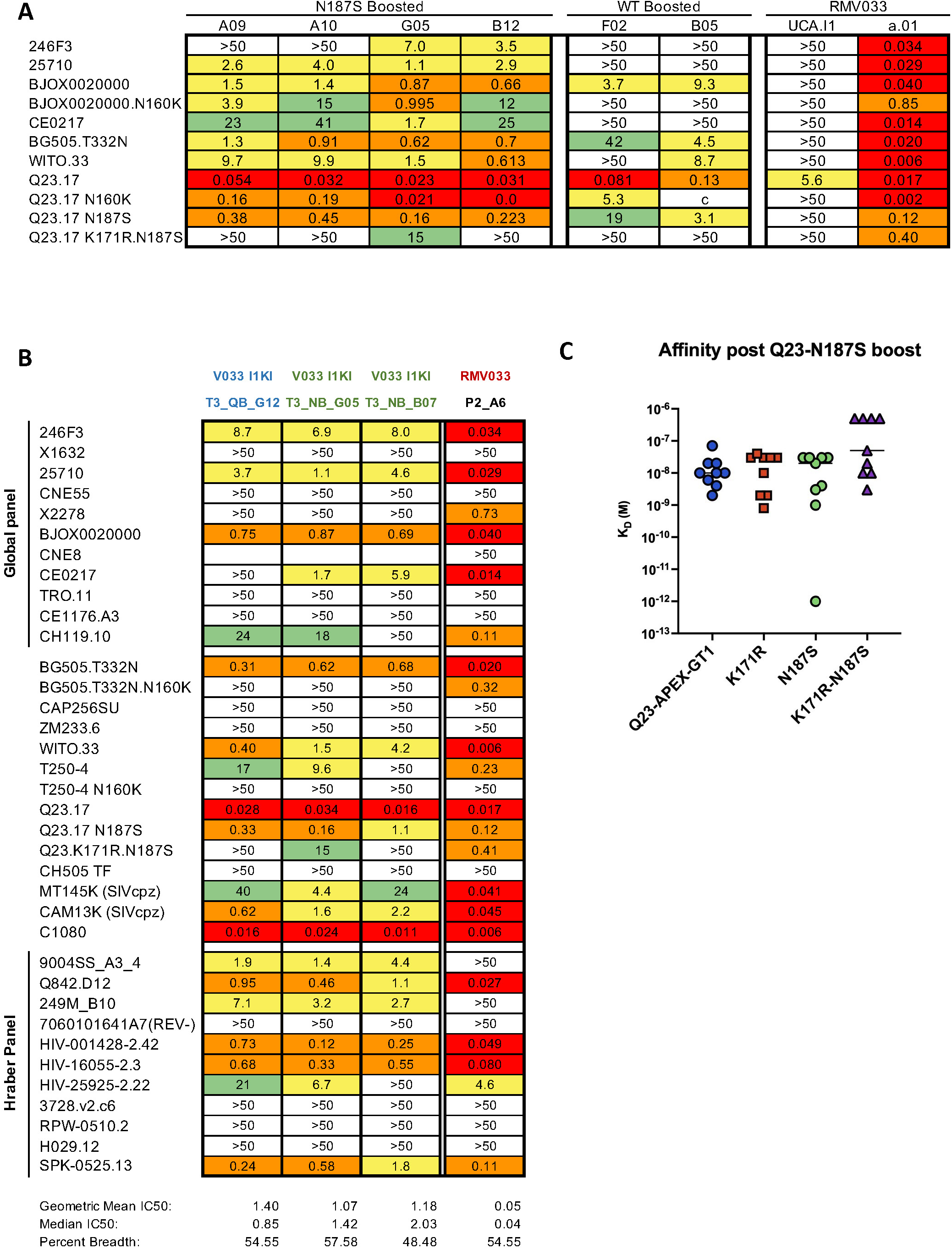
Neutralization breadth and potency of antibodies derived after homologous (Q23-APEX-GT1) or escape-variant (Q23-APEX-GT1 N187S) Env boost. (A) IC50 values of selected antibodies from 3 weeks post-boost and macaque mature V033 (RM V033 P2A6) against autologous (Q23.17), Q23.17 N160K and escape variants (Q23.N187S, Q23.K171R.N187S Q23.R169E) and limited panel of 5 heterologous tier-2 HIV-1 strains. (B) IC50 values of three antibodies against a 37-member panel of HIV-1 strains including 11 from the Tier-2 Global panel and 11 from the Hraber panel. Nomenclature: T = time in weeks post-boost; QB= Q23-APEX-GT1 Boost; NB= Q23-APEX-N187S Boost; final alphanumeric triplet = mAb identity (C) BLI binding values of boost-derived antibodies against autologous Q23 Env and escape-variant Envelopes.

To provide molecular characterization of boost-derived antibodies, we determined the cryo-EM structures of homologous-boosted T3_QB_G12 Fab and variant-boosted T3_NB_G05 Fab in complex with their respective boosting Env trimers (**Fig. 9, Figs. S12&S13, Supplemental Table 1**). The 3D reconstruction densities for the T3_QB_G12 and T3_NB_G05 Fab-trimer complexes extended to 3.8 Å and 3.1 Å resolution, respectively, and revealed both boost-derived antibodies to bind 3 Fabs per trimer like other V033-a lineage members and variants (**Fig. S12)** (Roark et al., 2024). While T3_NB_G05 recognized the V2-apex with an angle of approach similar to prime-derived T6_P_H03 and mature V033-a.01, T3_QB_G12 was modestly rotated and largely did not superimpose with the other Fab structures (**Fig. 9A**). As a result of this rotation, the T3_QB_G12 light chain was positioned in close proximity to adjacent protomer B and recognized the hypervariable V2 loop through LCDR1 (**Fig. 9B, left**). We propose that T3_QB_G12 is rotated to prevent clashes between N160 glycan and SHM Y53 in HCDR2, which is significantly larger than Gly or Ala residues present in all other Fabs and would provide stronger interactions by stacking against the first two *N*-acetylglucosamine residues of N160 glycan_A_ (**Fig. 9B, right, 9C**). Despite this rotation of the T3_QB_G12 Fab body relative to others, the extended HCDR3 tips of each antibody aligned at the C-strand, likely owing to the conformational restraints imposed by the V033-a-class defining antiparallel mainchain hydrogen bonds with the C-strand, which were maintained in the boost-derived antibodies (**Fig. 9A, Fig. S12).**

**Figure 9.**
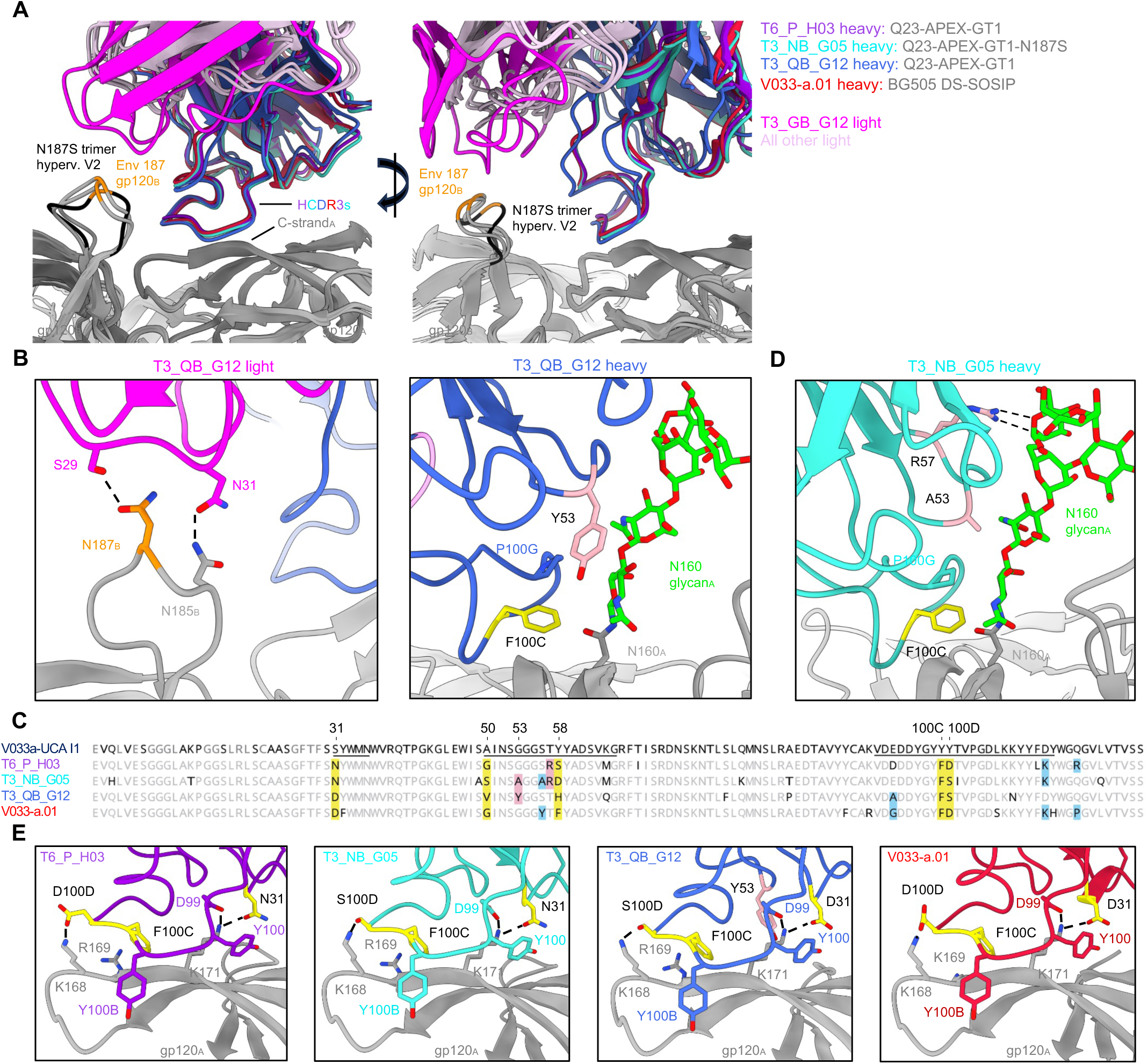
Structures of homologous and variant-derived antibodies reveal structural basis of heterologous neutralization breadth induced by boosting. (A) Orthogonal views for the gp120 alignment of T6_P_H03, T3_NB_G05, T3_QB_G12, and V033-a.01 cryo-EM structures in complex with Env SOSIP trimers. Only one Fab per trimer is shown for clarity, and HCDR3 secondary structure is also removed for clarity. The position of hypervariable V2 loop residue 187 is highlighted in orange on the neighboring protomer_B_ for each structure. This loop is colored black in the mutant N187S trimer in complex with T3_NB _G05 to highlight its distinct conformation. (B) Unique T3_QB _G12 interactions with Env. Left; The rotation of T3_QB_G12 positions its light chain proximal to hypervariable V2 loop on the adjacent protomer, resulting in interactions via LCDR1. T6_P_H03, T3_NB_G05, and V033-a.01 do not recognize this loop in their respective structures. Right; proposed explanation for the modest rotation of T3_QB_G12 relative to all other Fabs. The rotation of T3_QB _G12 Fab results from accommodation of the G53Y mutation in HCDR2 (pink) so it does not clash with N160 glycan on protomer_A_, which is significantly larger than Gly or Ala residues present in all other Fabs. Select paratope residues are shown in stick representation and are colored according to panel C if they are a result of somatic hypermutation. Residues are labeled by Kabat numbering. (C) Heavy chain amino acid sequences of T6_P_H03, T3_NB_G05, T3_QB_G12, and V033-a.01 are aligned to the reference V033a UCA I1. Residues matching the reference are depicted in light gray and nonmatching residues are depicted in black. The HCDRs are underlined in the I1 sequence. Conserved positions of somatic hypermutation shared by all four antibodies are highlighted in yellow and mutated positions identical to V033-a.01, but only appearing in one or two murine antibodies, are highlighted in light blue. The unique mutated residue Y53 in T3_QB_G12 indicated in panel B is highlighted in pink. (D) Unique T3_NB _G05 interactions with N160 glycan. Select paratope residues are shown in stick representation and are colored according to panel C if they are a result of somatic hypermutation. (E) Comparison of C-strand interactions for each Fab. Interacting residues are shown in stick representation and are colored yellow or pink according to the Top alignment. HCDR3 secondary structure is removed to facilitate visualization. Residues are labeled by Kabat numbering.

Similar to T3_QB_G12 Fab, T3_NB_G05 also acquired SHM within HCDR2 to increase interactions with N160 glycan_A_ (**Fig. 9C&D**). SHM residue A53 increases the hydrophobic interactive surface of the HCDR2 paratope over germline G53, while SHM R57 extends further than germline T53 and forms multiple hydrogen bonds with a terminal mannose residue. Boosting resulted in an increased frequency of antibodies bearing the improbable SHM residue K101 at the HCDR3 base, which is shared between V033-a.01, T3_NB_G05, and prime-derived antibody T6_P_H03; cryo-EM structures for each of these antibodies reveal K101 to interact with a terminal mannose residue on N156 glycan_A_ (**Fig. S12**). Despite the variations in glycan interactive surfaces for each Fab, conformations of apical glycans were nearly identical between V033-a.01 and each of the murine antibodies (**Fig. S12**). The remaining sites of heavy chain SHM within the boost-derived antibody paratopes closely mirrored the pattern of maturation described for prime-derived antibody T6_P_H03 and was conserved with the mature rhesus bnAb V033-a.01, resulting in stronger interactions with C-strand residues K168 and K171 through conserved structural mechanisms (**Fig. 9C, 9E**). T3_QB_G12 and T3_NB_G05 each acquired light chain SHM at just 2 of 7 identical positions acquired by V033-a.01, none of which fell within their respective interactive surfaces (**Fig. S12**). However, in the context of V033a-UCA I1, introduction of each of these light chain mutations individually improves neutralization of the autologous Q23.17 virus 2 to 3-fold (Habib et al., 2025), suggesting a role for stabilizing structural components of the Fab itself that may synergize with heavy chain maturation to yield the modest increases in breadth and potency of boost-derived antibodies.

Together, these data suggest a single prime followed by a lineage-based boost mirroring natural co-evolution can elicit and shepherd the V033a-UCA I1 toward V033-like bnAbs, with several murine variants achieving neutralization breadth equal to or greater than the mature V033-a.01 antibody via reproducible structural solutions for improved V2-apex recognition.

## Discussion

BnAb induction through vaccination remains a central goal for an effective HIV vaccine. Here, we leveraged a knockin mouse model expressing a macaque bnAb precursor (V033a-UCA I1), isolated following SHIV-Q23.17 infection, to demonstrate that a stabilized germline-targeted Env trimer immunogen—closely mimicking the native Env structure—can effectively engage macaque BCRs through immunization. Strikingly, this near-germline V033-a B cell precursor required only a short maturation pathway to develop into a bona fide bnAb in macaques. Consistent with this, a single immunization with adjuvanted Q23-APEX-GT1 protein trimer or Q23-APEX-GT1 saRNA encoding the membrane-bound trimer was sufficient to initiate on-track mutations in the V033a-UCA I1 lineage that closely mirrored those in the rhesus V033-a bnAb lineage but in a shorter timeframe and that correlated with the emergence of neutralization breadth. Furthermore, boosting with a Q23 Env lineage-based escape variant differing by only a single N187S mutation resulted in enhanced selection of these breadth-conferring on-track mutations and further acquisition of neutralization breadth and potency. Together, our data highlight that effective priming with a near native-like trimer may be the most critical step in eliciting V2 apex bnAbs. Whether that is true in humans, where the D gene repertoire is different, and probably less favorable, than in rhesus macaques, remains to be determined. Despite the lack of germline orthologue, human naïve repertoire can create similar EDDYG motif through addition of non-templated amino acids (Mishra et al., *Immunity* in review). Overall, we provide proof-of-concept that using a near-native HIV Env-derived germline-targeting priming immunogen alongside lineage-guided variants could serve as a promising vaccine strategy for bnAb induction.

Inferred germline (iGL) or bnAb precursor knockins have been tremendously helpful in testing immunogens that target very rare human precursors (Dosenovic et al., 2015; Huang et al., 2020; Jardine et al., 2015; McGuire et al., 2016; Steichen et al., 2019). While knocking in unrearranged human immunoglobulin loci to allow for recombination is an attractive option (Luo et al., 2023; Tian et al., 2016), the low number of total B cells in a mouse can make targeting desired precursor B cells extremely difficult (Sok et al., 2016). Knocking in a pre-rearranged BCR, in contrast, allows for a controlled titration of precursor frequency through adoptive transfer, as well as the opportunity to observe the potential recapitulation of known bnAb developmental pathways (Abbott et al., 2018; Lin et al., 2018; Steichen et al., 2019; Wang et al., 2021). While xenografted “macaquized mice” have been used in the HIV field to study viral amplification (Metcalf Pate et al., 2015), to our knowledge the V033a-UCA I1 is the first immunoglobulin knockin mouse to bridge the results from these two primary animal models for preclinical HIV-1 vaccinology.

Compared to bnAbs targeting epitopes in HIV-1 Env such as CD4bs, V2-apex targeting bnAbs such as V033 do not require very high levels of SHM, but they do require a long HCDR3 loop (generated via rare V-D-J recombination events) with a preference for C-strand anionic CDRH3 residues (Andrabi et al., 2015; Habib et al., 2025; Roark et al., 2024). We previously demonstrated that certain HIV Envs containing a naturally occurring glycan-sparse apex can readily bind to unmutated precursor versions of V2-apex bnAbs when presented in a near-native-like Env trimer configuration (Andrabi et al., 2015; Gorman et al., 2016; Voss et al., 2017). Using macaque SHIV infection carrying HIV-1 T/F envelopes and other glycan-sparse Env, our group found that particular Env properties select for classes of V2 apex bnAb precursor which share binding approaches with human V2-apex targeting bnAbs (Habib et al., 2025; Roark et al., 2024). In one of those studies, we also observed that certain envelopes, such as Q23.17, have a higher propensity to induce V2 apex targeting bnAbs than other Envs (Habib et al., 2025). Notably, SHIV-Q23.17 rapidly elicited the rhesus V033-a V2 apex bnAb lineage in a SHIV-infected monkey. Lineage tracing during macaque SHIV infection, therefore, informed both the B cell lineage targeting and the Env selection for immunogen design in this study.

Germline targeting using stabilized versions of near-native Envs can offer a simplified approach to priming and boosting V2 apex bnAbs. A distinguishing feature of this approach is that once primed, early bnAb precursors can rapidly affinity-mature to a native-like Env as opposed to one more heavily mutated. In the V033 monkey, the neutralization potency of maturing bnAb lineage members to viruses expressing Q23.17 Env increased by >50,000 fold (Habib et al., 2025). In the knockin mice, maturing bnAbs increased their potency to Q23.17 virus. Coincident with this remarkable enhancement of potency to infecting autologous virus strain was the development of neutralization breadth. Faithful preservation of structural and antigenic features of an epitope on target Env glycoproteins is essential to generate protective antibody responses (Graham et al., 2019). To design the Q23 Env lineage-based immunogens for evaluation in the V033a-UCA I1 knockin mouse model, we utilized antibody-guided structure-based design to generate a prefusion-stabilized, native-like Q23 HIV Env trimer—Q23-APEX-GT1—with enhanced capacity to engage V2-apex bnAb precursors. In the absence of SHIV infection, which provides a constant source of evolving immunogen, escalating or interval immunizations have been shown to sustain GCs for longer duration (Bhagchandani et al., 2024). Here, however, we showed that a single bolus immunization of Q23-APEX-GT1 was sufficient to sustain GCs and allow V033 precursors to acquire substantial SHM, breadth, and potency, demonstrating that maturation into a bnAb does not always require multiple boosting immunizations. Although longitudinal analysis of bnAb development has been done only for very few human bnAb lineages, human bnAb lineages at intermediate stages of development (∼34 weeks) have been shown to neutralize autologous infective viruses (Bhiman et al., 2015). The experimental vaccinology here supports the notion that B cell precursors targeting the V2-apex epitope can be activated by native-like trimers without the need for extensive trimer modifications specifically designed to enhance precursor engagement.

The rapid maturation of macaque precursor V033a-UCA I1 in the absence of a replicating virus with a single immunization highlights the possibility that the features present in the early V033 lineage present a low barrier to developing into a bnAb. Notably, the rhesus line is characterized by the use of a D gene (D3-15*01) which encodes an ‘EDDYG motif’ similar to the germline encoded YYD motif in several human V2-apex bnAbs, likely aiding in the development towards the bnAb (Habib et al., 2025). While these germline-encoded EDDYG motifs are not uncommon in humans (Habib et al., 2024) (Mishra et al., *Immunity* in review), their potential contribution in human bnAb development remains underexplored. Furthermore, V033 is a less common V2-apex targeting antibody, in that it is trimer-preferring rather than trimer-specific. The binding of three Fabs to one trimer, as observed in the cryo-EM structure here, can enhance tensile strength during antigen extraction due to higher avidity during immunological synapse formation (Nowosad et al., 2016). Targeting similar precursors in humans may provide a route to a more tractable vaccination regimen.

Overall, our study provides proof-of-concept that a single immunization with a stabilized, near-native Q23-APEX-GT1 trimer can rapidly mature V2 apex bnAb precursors into functional antibodies, highlighting a promising vaccine development strategy for HIV-1.

## Supporting information

All Supplemental Figures and Tables

## Acknowledgements

We would like to thank all members of the Ragon Institute’s Batista lab and Flow Cytometry Core, as well as Anastasia Yandulskaya-Blue at the Scientific Editing Platform.

## Author Contributions

A.R.G. characterized the knockin mice, performed in vivo and 10X experiments, analyzed the data and drafted the manuscript. A.R.G., N.M., R.H., selected and expressed monoclonal antibodies. N.M., S.C., D.R.B, and R.A. conceived and designed experiments. R.S.R performed cryo-EM. M.A. and T.P. contributed animal experiments. A.A.A. assisted in cell sorting. J.D.A., N.E.J., and M.C. performed glycan analysis. N.M., S.C., G.A., Xu.L., B.L., K.A., P.O., R.R.C., and R.N. designed and prepared immunogens, performed neutralization and binding assays. N.M., S.C., G.A., Xu.L., B.L., tested monoclonal antibodies and trimers and probes for B cell sorting. J.E.W., M.B., and P.M. contributed to 10x sequencing experiments and analysis. J.R.E.P., M.S., L.X., and U.N. generated the V033a UCA I1 mouse. S.R.W. contributed to manuscript drafting and editing. X.L. oversaw lab infrastructure and mouse line maintenance. J.T. and A.B.W. generated and analyzed negative stained electron microscopy data. Y.P. and B.H.H. provided reagents. L.M. coordinated material needs, performed replicon design, synthesis and QC oversight, LNP prep and QC oversight, SMNP prep and QC oversight. N.C. performed LNP synthesis and QC. J.D. and L.R. performed replicon synthesis and QC. A.W. performed SMNP synthesis and QC. F.D.B. conceived of, designed and oversaw murine immunocharacterization experiments.

## Data and Materials Availability

BCR sequences and structural PDBs will be submitted to databases on acceptance and made publicly available upon publication and are available to reviewers upon request. Model animals available from corresponding author FDB on request and subject to standard MTA with Mass General Hospital.

## Materials and Methods

### Immunogen Design

To develop prefusion-stabilized, native-like Q23 Env trimers, a structure-guided engineering strategy was employed. Initial designs combined elements from previously described native flexibly linked (NFL) and repair-and-stabilization (RnS) scaffolds (Rutten et al., 2018; Sharma et al., 2015), supplemented with additional gp120 and gp41 stabilizing mutations targeting conformational hotspots. Mutations were introduced to enhance apex hydrophobic packing, restrict CD4-induced rearrangements, and minimize exposure of non-neutralizing V3 epitopes. Constructs were designed as covalently linked single-chain trimers (SCTs) using a glycine-serine linker between gp120 and gp41. Core stabilizing mutations— including SOSIP (501C–605C, I559P) and a 201C–433C disulfide bond—were retained across all ten designs (Q23-SCT21 to Q23-SCT30). Subsequent variants introduced glycine substitutions (positions 569, 636), rigidity-enhancing mutations (Q551P), fusion peptide modifications (519R–520R), and RnS-derived residues to reinforce gp41. For gp120, select mutations (49E, 153E, 219A, 302Y, 320M, 334S) were incorporated based on structural analysis and computational modeling (ADROITrimer pipeline). Synthetic genes encoding the amino acid sequences for SCT21-30 were ordered as gBlocks and cloned in phCMV3 vector in-frame with tPA signal peptide with Gibson assembly.

### Stabilized Env Expression and Purification

Stabilized HIV Env trimers were expressed in HEK293F cells transfected with Env-encoding plasmids using PEI-MAX 4000 (Polysciences, Inc.). Four days post-transfection, supernatants were harvested and trimers purified via affinity chromatography using either GNL-agarose or PGT145-conjugated CNBr-activated Sepharose 4B beads (GE Healthcare). Eluates were further purified by size-exclusion chromatography (SEC) on a Superdex 200 10/300 GL column (GE Healthcare) and eluted in PBS or TBS.

### Biolayer Interferometry

For antigenic screening of Q23 SCT immunogens, BLI was performed with purified monoclonal antibodies (10 µg/ml). For assessing the polyclonal serum IgG response in immunized animals, purified polyclonal serum IgG (10 µg/ml) was used. IgGs were immobilized on ProA sensors (Sartorius) to a signal of 1.0 nM using an Octet Red96 instrument (ForteBio). The immobilized IgGs were then dipped in the running buffer (PBS, 0.1% BSA, 0.02% Tween20, pH 7.4) followed by 500 nm of redesigned Q23 trimers, or running buffer. Following a 120 s association period, the tips were dipped into the running buffer and dissociation was measured for 240 s.

### Cell Surface Binding Assay

HEK293T cells were seeded in T75 flasks and transfected with plasmids encoding the antigen of interest using Lipofectamine 2000 (Thermo Fisher Scientific, Cat. #11668500), according to the manufacturer’s instructions. Forty-eight hours post-transfection, cells were harvested using FACS buffer (PBS supplemented with 2% FBS and 5 mM EDTA; Invitrogen, Cat. #15575-038), washed twice, and resuspended at 1 × 10⁶ cells/mL.

Cells were incubated with primary antibodies of interest at a final concentration of 10 µg/mL for 1 hour at 4°C. After three washes with FACS buffer, cells were stained with PE-conjugated mouse anti-human IgG Fc secondary antibody (SouthernBiotech, Cat. #9040-09) for 1 hour at 4°C in the dark. Following additional washes, cells were resuspended in FACS buffer and analyzed on a Bio-Rad ZE5 flow cytometer. Data were processed using FlowJo software (BD Biosciences).

### Replicon RNA Synthesis and LNP Formulation

Env trimer coding sequences were cloned into a VEE-based replicon backbone. For replicon RNA synthesis, plasmid constructs were linearized via endonuclease digestion and purified with PureLink PCR Purification columns (ThermoFisher). Linearized plasmid DNA was transcribed via *in vitro* transcription using HiScribe T7 High Yield RNA Synthesis Kit (NEB), purified using PureLink RNA Mini columns (ThermoFisher), post-transcriptionally capped and methylated using ScriptCap Cap 1 Capping System (CellScript), and purified again using RNA Mini columns. To encapsulate replicons in LNPs, lipids including Heptadecan-9-yl 8-((2-hydroxyethyl)(6-oxo-6-(undecyloxy)hexyl)amino)octanoate (SM-102, Broadpharm), 1,2-dioctadecanoyl-sn-glycero-3-phosphocholine (DSPC, Avanti Polar Lipids), cholesterol (Chol, Avanti Polar Lipids), and 1,2-dimyristoyl-rac-glycero-3-methoxypolyethylene glycol-2000 (PEG-DMG, Avanti Polar Lipids) were dissolved in ethanol at a 50:10:38.5:1.5 SM102:DSPC:Chol:PEG-DMG molar ratio. RNA was dissolved in 10 mM citrate buffer pH 3, and then combined with the lipid solution at an N:P ratio of 6:1 (N, number of nitrogens on SM-102 ionizable lipid; P, number of phosphate groups on RNA) using an Ignite microfluidic mixing system (Precision Nanosystems) at a flow rate of 12 ml/min and a lipid:mRNA volume ratio of 3:1. The resulting replicon-loaded LNPs were dialyzed in 20 mM Tris-acetate and 8% (w/v) RNAse-free sucrose (VWR) using 3500 MWCO Slide-A-Lyzer cassettes (ThermoFisher) and stored at −80 °C till use.

SMNP adjuvant was prepared as previously described (Silva et al., 2021).

### Animals and Immunizations

10–12-week-old CD45.1^+/+^ mice (B6.SJL-Ptprc^a^ Pepc^b^/BoyJ) were purchased from the Jackson Laboratory (Bar Harbor ME). Mice with B cells expressing a macaquized V033A-UCA I1 BCR were generated using the same CRISPR/Cas9 approach previously described for human knockins (Lin et al., 2018; Wang et al., 2021). Mice were housed in a 12-hr light-dark cycle with ad libitum access to water and standard laboratory chow. All animal procedures were conducted in accordance with protocols approved by the Institutional Animal Care and Use Committee (IACUC) at Mass General and conducted in accordance with the regulations of the Association for Assessment and Accreditation of Laboratory Animal Care (AAALAC) International. B cells from donor V033a-UCA I1 IgH^+/WT^ IgL^+/WT^ were isolated using a Pan B cell isolation kit II (Miltenyi Biotec, Bergisch Gladbach, Germany) and were adoptively transferred through tail vein injection into acceptor B6.SJL-Ptprca Pepcb/BoyJ (CD45.1 mice) at frequencies indicated in the text. Mice were immunized 24 hours post adoptive transfer. Groups of mice were injected bilaterally in the hind leg with Q23-APEX-GT1 LNPs at 2 µg total replicon dose (1 ug per injection site) or subcutaneously at the base of tail with 10 µg protein trimer antigen and 5 µg SMNP total (administered bilaterally with dose split equally between the two sites).

### Post-Immunization Analysis

For animals receiving SC immunization draining inguinal lymph nodes and for animals receiving IM immunization iliac and popliteal lymph nodes were harvested from mice at weeks 2, 4, 6 post-immunization, as described in related figure legends. Single-cell suspensions were prepared by gently crushing the draining lymph nodes from a single mouse together and passing them through a 70 μM strainer. Incubation with PBS containing Live/Dead Blue (Thermo Scientific, Waltham MA) diluted 500-fold and FcR Blocking reagent (Purified Rat anti-mouse CD16/CD32, BD Biosciences) diluted 200-fold was done for 20 mins at 4°C. After washing, BCR antigen staining was performed using biotinylated Q23-APEX-GT1 trimer conjugated to either streptavidin-BV510 (BioLegend, San Diego CA) or strepatvidin-Alexa647 (BioLegend) for 30 mins at 4°C. Excess antigen was washed off and cell surface staining was performed with an antibody cocktail containing CD4, CD8, F4/80, GR-1, NK1.1 (APC-eFluor 780, eBioscience, clone RM4-5, 53-6.7, BM8, RB6-8C5, PK136 respectively), B220 (BUV395, BD Bioscience, clone RA3-6B2), CD38 (BUV563, BD Biosciences, clone 90), CD95 (PE-Cy7, BioLegend, clone L138D7), CD45.1 (BV605, BioLegend, clone A20), CD45.2 (PE, BD Biosciences, clone 104), CD138 (BV650, BD Biosciences, clone 281-2), IgD (Alexa 594, Biolegend, 11-26c.2a clone) and IgM (BV750, II/41 clone) for 30 mins at 4°C. Flow cytometry data was acquired using BD FACS Symphony A5 cell analyzer. For cell sorting, Live/Dead stain was replaced with SYTOX Green (Thermo Fisher Scientific)/DAPI. The antibodies used for sorting were CD4, CD8, F4/80, GR-1 & NK1.1 (APC-eFluor 780, eBioscience, clone RM4-5, 53-6.7, BM8, RB6-8C5, PK136 respectively), B220 (Alex Fluor594 BioLegend, clone RA3-6B2), CD38 (BB700, BD Biosciences, clone 90), CD95 (PE-Cy7, BioLegend, clone L138D7), CD45.1 (APC R700, BD Biosciences, clone A20), CD45.2 (PE, BD Biosciences, clone 104), IgD (BV605, BioLegend, clone 11-26c.2a), CD138 (BV650, BD Biosciences, clone 281-2). Cells from each individual mouse were barcoded with TotalSeq™-C anti-mouse Hashtag Antibodies. A total of 10 hashtags were used. Cells were washed 3 times to remove any excess antibodies.

### Cell Sorting and Paired BCR Sequencing

Cells were sorted using BD FACSymphony S6 (BD Franklin Lakes, NJ) using 85 μM nozzle. Samples were sorted into PCR tubes containing PBS with 10% FBS. Encapsulation of sorted cells and NGS library preparation was performed following the 10x Genomics Chromium Next GEM Single Cell 5’ Reagent Kits v2 protocol (10x Genomics). TapeStation Systems D5000 high sensitivity Screen Tape assay (Agilent, Santa Clara, CA) was used to measure library size. After quantifying the libraries through Qubit dsDNA High Sensitivity (Invitrogen, Waltham MA), they were pooled and were run on NextSeq 550 System (Illumina, San Diego, CA). Analysis was performed using the Cell Ranger v.6 software pipeline (10x Genomics, Pleasanton, CA) with a customized reference database. Sequencing data was analyzed using Geneious Prime and Biologics software (Geneious, Auckland, New Zealand) and IMGT/V-Quest (Brochet et al., 2008; Giudicelli et al., 2011, 2022).

### Monoclonal Antibody Expression and Purification

Paired heavy and light chain plasmids for monoclonal antibodies cloned from immunized animals were co-transfected in Expi293 cells (Thermo Fisher Scientific, Waltham MA) in a 1:1 ratio using FectoPRO transfection reagent. Monoclonal IgGs were purified from the culture supernatant five days post-transfection with Protein-A Sepharose beads (Cytiva, Marlborough MA) per manufacturer’s instructions. After elution with IgG elution buffer (Thermo Fisher Scientific), antibodies were buffer exchanged into Tris buffered saline (TBS).

### Monoclonal Fab preparation for Cryo-EM structural studies

Monoclonal IgG Fab heavy chain constructs were engineered by inserting a His-Avi tag followed by a stop codon upstream of the disulfide bond in the Fc region. Paired plasmids encoding the truncated heavy and light chains were co-transfected into Expi293 cells (Thermo Fisher Scientific, Cat# A14527) at a 1:1 ratio using FectoPRO transfection reagent (Polyplus, Cat# 116-001). At 24 hours post-transfection, cells were supplemented with 0.3 M valproic acid (Sigma, Cat# P4543-100G) and 40% glucose (Gibco, Cat# A2494001). Five days after transfection, Fabs were purified from the culture supernatant using Ni Sepharose 6 Fast Flow resin (Cytiva, Cat# 17531802) following the manufacturer’s protocol. Eluted Fabs were buffer-exchanged into PBS and concentrated using a 10 kDa molecular weight cutoff centrifugal filter unit (Millipore, Cat# UFC905024). The concentrated Fab proteins were further purified by size exclusion chromatography on a Superdex 200 Increase 10/300 GL column (Sigma-Aldrich, Cat# GE28-9909-44). Fractions corresponding to the monomeric Fab peak were pooled, reconcentrated, and used for downstream Cryo-EM studies.

### Site-specific Glycosylation Analysis

To analyze glycosylation of the Q23_SCT27_GT2.V1 trimer, 100 µg of protein was denatured in 50 mM Tris/HCl (pH 8.0) with 6 M urea and 5 mM DTT for 1 hour, then alkylated with 20 mM iodoacetamide (IAA) in the dark for 1 hour. Residual IAA was quenched with 20 mM DTT. After buffer exchange into 50 mM Tris/HCl, the sample was split into three ∼33 µg aliquots for separate overnight digestions with Trypsin, Chymotrypsin, or Alpha-lytic protease (1:30 w/w) at 37 °C. Peptides were desalted using Oasis HLB µElution plates and analyzed by nanoLC-ESI-MS using an Easy nLC 1200 coupled to an Orbitrap Eclipse with stepped HCD. Peptides were separated on an EasySpray PepMap RSLC C18 column with a 280-minute gradient. Key MS settings included a scan range of 300−2000 m/z, HCD energies of 15/25/45%, MS1 resolution of 120,000, and MS2 at 30,000. Glycopeptide data were processed in Byos (v5.5). True positives were confirmed by oxonium ions and matching b/y fragments. Searches used the Protein Metrics 305 N-glycan library with added sulfated glycans. Glycan abundance was calculated by chromatographic area comparison of glycoforms with identical peptide sequences. A 1% FDR was applied (4 ppm precursor, 10 ppm fragments), and all charge states were summed. Glycans were classified by composition: HexNAc(2)Hex(9–3) as M9–M3; HexNAc(2)Hex(10+) as M9Glc; fucosylated variants as FM; Hybrid types included HexNAc(3)Hex(5–6)X and Fhybrid; complex types were based on HexNAc count and fucosylation. Core glycans were <M3, and M9Glc–M4 were oligomannose. High-mannose included both oligomannose and hybrid types.

### Neutralization Assays

Neutralizing antibody assays were performed using TZM-bl indicator cells, as previously described (Roark et al., 2021). Briefly, TZM-bl cells were grown in cell culture medium (DMEM supplemented with 10% FBS and 1% penicillin/streptomycin), were plated in 96 well plates at a concentration of 1.5×10^4^ cells/well and incubated at 37°C overnight. Serum samples were heat inactivated at 56°C for one hour, and serial 5-fold dilutions, starting at a dilution of 1:100, were made in cell culture media supplemented with 1% normal mouse serum (Avantor, Cat# 76226-474; supplied by Rockland Immunochemicals, original item# D108-00-0100). Normal mouse serum was included in the diluent to keep the concentration of test serum constant across wells. For monoclonal antibody neutralization assays, mAbs were serially diluted starting at a concentration of 50 µg/ml in cell culture media lacking normal mouse serum. Viruses were diluted to achieve a multiplicity of infection (MOI) of 0.3 upon addition to TZM-bl cells. Viruses were incubated with serum or antibody dilutions at 37°C for one hour, after which the virus-serum or virus-antibody mixtures were plated onto adherent TZM-bl cells and incubated at 37°C for 48 hours. Following incubation, cells were lysed with 0.5% Triton-X 100 in PBS and luciferase activity levels measured using the Promega luciferase assay system (Promega Cat# E1501) on a BioTek Synergy Neo2 plate reader (Agilent Technologies). Neutralization IC_50_ and ID_50_ values were calculated using Prism 10 software.

### Cryo-EM sample preparation and data collection

The structures of murine V033 variant Fabs in complex with envelope trimer were determined using single-particle cryo-EM. Samples were prepared by mixing each Fab with the cognate trimer used for terminal immunizations (T6_P_H03 with Q23-APEX-GT1; T3_NB_G05 with Q23-APEX-GT1.N187S; and T3_QB_HG12 with Q23-APEX-GT1) at a 2:1 Fab:protomer ratio and incubating on ice for 30 minutes. Each sample was then supplemented with PBS and the detergent Dodecyl β-D-maltoside (DDM; Anatrace) to achieve a final trimer concentration of ∼2.5 mg/mL and final DDM concentration of 0.005% (w/v). 3 uL of each sample was then applied to a freshly glow-discharged (20 seconds at 20 mA) copper C-flat Holey carbon-coated grid (CF-1.2/1.3 300 mesh; EMS) within the chamber of a Vitrobot Mark IV at room temperature with 100% humidity. After incubating for 30 seconds, grids were blotted for 3 seconds prior to immediate plunge freezing in liquid ethane. Single-particle cryo-EM data were collected on a FEI Titan Krios cryo-transmission electron microscope operating at 300 kV and equipped with a Gatan K3 detector. Automated data collection was carried out using Leginon (Suloway et al., 2005) in counting mode with a 105,000x magnification and a pixel size of 0.83 Å. The total dose of 58 e^-^/Å^2^ was fractionated over 50 raw frames, with defocus values set to cycle between −0.80 and −2.0 μm. Initial data collection attempts revealed prohibitive preferred orientation of the 3:1 Fab:trimer-bound complexes, therefore a 30-degree tilt was applied during collection of the final datasets used for each of the respective reconstructions.

### Cryo-EM data processing and model building

All processing was done in cryoSPARC v3.4 (Punjani et al., 2017), including micrograph curation, motion correction, CTF estimation, particle picking, 2D classification, *ab initio* modeling, and 3D refinements. Particle picking was performed with non-templated blob picker. Multiple rounds of 2D classification, followed by iterative *ab initio* modeling and heterogenous refinements, were used to curate the particle datasets used in the final 3D reconstructions for each sample. All *ab initio* modeling and homogenous 3D refinements were performed using C1 symmetry, while all non-uniform 3D refinements of were performed using C3 symmetry. The initial coordinates for each murine V033 variant complex were obtained by docking the V033-a.01 Fab from PDB-9BNP and the Q23-APEX-GT2 trimer from PDB-9NVV into the present cryo-EM density using UCSF ChimeraX (Pettersen et al., 2021). Sequence corrections from the initial models to match each respective sample Fab and trimer were done manually in Coot (Emsley & Cowtan, 2004). Each of the atomic models were solved by iterative real-space refinement in Phenix (Adams et al., 2004) and manual rebuilding in Coot. Overall structure quality was assessed using MolProbity (Davis et al., 2004) and EMRinger (Barad et al., 2015). Structural superimpositions of complexes were performed in UCSF ChimeraX. Final model statistics and validations are provided in **Fig. S13** and **Supplementary Table 1.**

