## Supplementary material for "Rapid acquisition of HIV-1 neutralization breadth in a rhesus V2 apex germline antibody mouse model after a single bolus immunization": All Supplemental Figures and Tables

### SUPPLEMENTAL MATERIALS

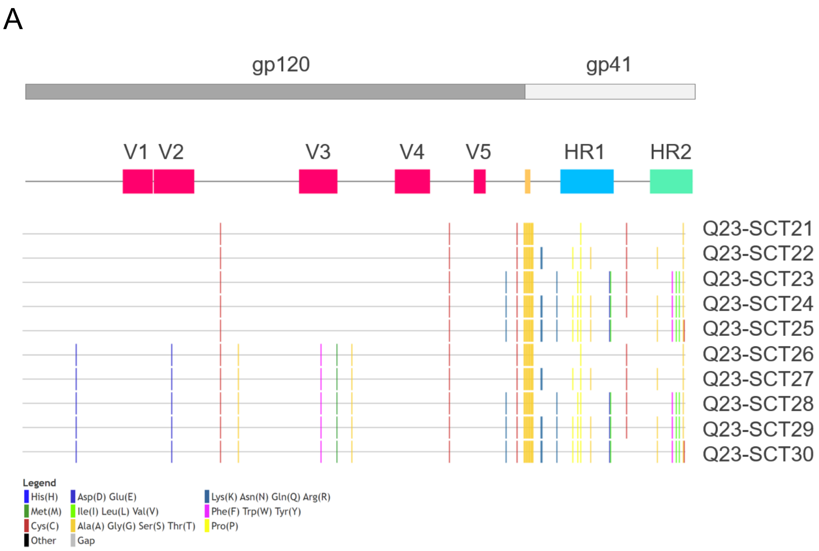


B

| Construct | Yield (mg) |
|---|---|
| Q23-SCT21 | 0.89 |
| Q23-SCT22 | 2.31 |
| Q23-SCT23 | 1.30 |
| Q23-SCT24 | 0.81 |
| Q23-SCT25 | 1.67 |
| Q23-SCT26 | 0.47 |
| **Q23-SCT27** | **2.02** |
| Q23-SCT28 | 1.72 |
| Q23-SCT29 | 2.16 |
| Q23-SCT30 | 2.46 |

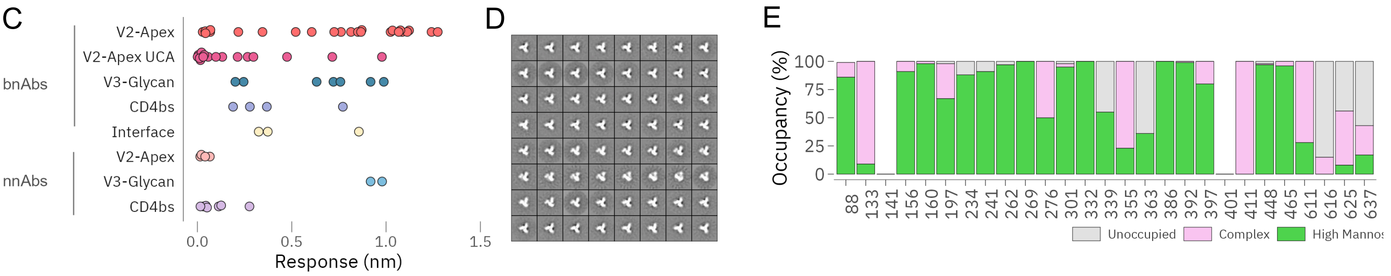

**Supplementary Figure 1. Further characterization of Q23-SCT immunogens, related to Figure 1.**

(A) Highlighter plots showing the mutations across all 10 SCTs compared to wildtype Q23 sequence. First box shows the placement of gp120 (colored dark gray) and gp41 (colored light gray) while the second box shows the placement of variable loops and the refolding regions of HR1-2 (Heptad Repeat 1-2). (G4S)2 linker is shown in yellow. Mutations are marked by colored vertical lines.

(B) Yield (in mg) of GNL-purified proteins prior to SEC purification from 100 ml of 293F. Q23-SCT27 is highlighted in bold.

(C) BLI binding responses of a large panel of 70 mAbs against all epitope class of HIV-1 Env trimer against Q23-SCT27. mAbs are segregated in two bins of bnAbs and non-nAbs.

(D) Negatively stained EM of Q23-SCT27 trimers with 2D-averaged classes.

(E) Proteomics-based site-specific glycan analysis (SSGA) of Q23-SCT27. High mannose, green; complex glycan, pink; unoccupied, gray. No signal could be resolved at PNGS 141 and 401.

```
Q23-WT             MDAMKRGLCCVLLLCGAVFVSPSQEIHARFRRGARAENLWVTVYYGVPVWRDADTTLFCASDAKAYETEKHNVWATHACVPTDPNPQEIHLDNVTEKFNMWKNNMVEQMHTDIISLWDQS
Q23-SCT21          ........................................................................................................................
Q23-SCT22          ........................................................................................................................
Q23-SCT23          ........................................................................................................................
Q23-SCT24          ........................................................................................................................
Q23-SCT25          ........................................................................................................................
Q23-SCT26          .....................................................E..................................................................
Q23-SCT27          .....................................................E..................................................................
Q23-SCT28          .....................................................E..................................................................
Q23-SCT29          .....................................................E..................................................................
Q23-SCT30          .....................................................E..................................................................
Clustal Consensus  *****************************************************:******************************************************************

Q23-WT             LKPCVKLTPLCVTLHCTNVTSVNTTGDREGLKNCSFNMTTELRDKRQKVYSLFYRLDIVPINENQGSEYRLINCNTSAITQACPKVSFEPIPIHYCTPAGFAILKCKDEGFNGTGLCKNV
Q23-SCT21          ..............................................................................C.........................................
Q23-SCT22          ..............................................................................C.........................................
Q23-SCT23          ..............................................................................C.........................................
Q23-SCT24          ..............................................................................C.........................................
Q23-SCT25          ..............................................................................C.........................................
Q23-SCT26          .............................E................................................C.................A.......................
Q23-SCT27          .............................E................................................C.................A.......................
Q23-SCT28          .............................E................................................C.................A.......................
Q23-SCT29          .............................E................................................C.................A.......................
Q23-SCT30          .............................E................................................C.................A.......................
Clustal Consensus  ***************************** ************************************************ *****************:***********************

Q23-WT             STVQCTHGIKPVVSTQLLLNGSLAEKNITIRSENITNNAKIIIVQLVQPVTIKCIRPNNNTRKSIRIGPGQAFYATGDIIGDIRQAHCNVTRSRWNKTLQEVAEKLRTYFGNKTIIFANS
Q23-SCT21          ........................................................................................................................
Q23-SCT22          ........................................................................................................................
Q23-SCT23          ........................................................................................................................
Q23-SCT24          ........................................................................................................................
Q23-SCT25          ........................................................................................................................
Q23-SCT26          ...........................................................Y...............M..............S.............................
Q23-SCT27          ...........................................................Y...............M..............S.............................
Q23-SCT28          ...........................................................Y...............M..............S.............................
Q23-SCT29          ...........................................................Y...............M..............S.............................
Q23-SCT30          ...........................................................Y...............M..............S.............................
Clustal Consensus  *********************************************************** *************** **************:*****************************

Q23-WT             SGGDLEITTHSFNCGGEFFYCNTSGLFNSTWYVNSTWNDTDSTQESNDTITLPCRIKQIINMWQRAGQAMYAPPIPGVIKCESNITGLLLTRDGGKDNNVNETFRPGGGDMRDNWRSELY
Q23-SCT21          ....................................................................C...................................................
Q23-SCT22          ....................................................................C...................................................
Q23-SCT23          ....................................................................C...................................................
Q23-SCT24          ....................................................................C...................................................
Q23-SCT25          ....................................................................C...................................................
Q23-SCT26          ....................................................................C...................................................
Q23-SCT27          ....................................................................C...................................................
Q23-SCT28          ....................................................................C...................................................
Q23-SCT29          ....................................................................C...................................................
Q23-SCT30          ....................................................................C...................................................
Clustal Consensus  ********************************************************************.***************************************************

Q23-WT             KYKVVEIEPLGVAPTRAKRRVVEREK------RAVGIGAVFLGFLGAAGSTMGAASITLTVQARQLLSGIVQQQNNLLRAIEAQQHLLKLTVWGIKQLQARVLAVERYLRDQQLLGIWGC
Q23-SCT21          ................C......GGGGSGGGGS...............................................P.......................................
Q23-SCT22          ................C......GGGGSGGGGS.......RR..............................P.......P.........G.............................
Q23-SCT23          .....K..........C......GGGGSGGGGS.......................N....................P..P............................EV.........
Q23-SCT24          .....K..........C......GGGGSGGGGS.......RR..............N...............P....P..P.........G..................EV.........
Q23-SCT25          .....K..........C......GGGGSGGGGS.......RR..............N...............P....P..P.........G..................EV.........
Q23-SCT26          ................C......GGGGSGGGGS...............................................P.......................................
Q23-SCT27          ................C......GGGGSGGGGS.......RR..............................P.......P.........G.............................
Q23-SCT28          .....K..........C......GGGGSGGGGS.......................N....................P..P............................EV.........
Q23-SCT29          .....K..........C......GGGGSGGGGS.......RR..............N...............P....P..P.........G..................EV.........
Q23-SCT30          .....K..........C......GGGGSGGGGS.......RR..............N...............P....P..P.........G..................EV.........
Clustal Consensus  *****:**********.******          *******  ************** *************** **** ** ********* ******************. *********

Q23-WT             SGKLICTTNVPWNSSWSNKSLDEIWNNMTWLQWDKEINNYTQLIYRLIEESQNQQEKNEKELLELD
Q23-SCT21          ......C........................................................A..
Q23-SCT22          ......C..............................G.........................A..
Q23-SCT23          ......C.............................................F...I..V...A..
Q23-SCT24          ......C..............................G..............F...I..V...A..
Q23-SCT25          .....................................G..............F...I..V...AC.
Q23-SCT26          ......C........................................................A..
Q23-SCT27          ......C..............................G................K........A..
Q23-SCT28          ......C.............................................F...I..V...A..
Q23-SCT29          ......C..............................G..............F...I..V...A..
Q23-SCT30          .....................................G..............F...I..V...AC.
Clustal Consensus  ****** ******************************.************** *** ** ***  *
```

**Supplementary Figure 2. Construct sequences, related to Figure 1.**

Alignment of all designed constructs with individual mutations relative to wildtype Q23 envelope glycoprotein sequence. Mutations are colored according to side-chain chemistry. Unchanged positions are shown by dots.

A

| Class | | mAb | IC50 | BLI Response | Surface |
|---|---|---|---|---|---|
| UCA | V2-Apex UCA | 42056-a.UCA | 10 | 0.01 | 931 |
| | | 40591-a.UCA | 10 | 0.02 | 1541 |
| | | RHA1 UCA | 10 | 0.21 | 3272 |
| | | 6561-a.UCA | 10 | 0.00 | ND |
| | | 6070-a.UCA | 10 | 0.27 | 1356 |
| | | T646-a.UCA | 10 | 0.10 | 961 |
| | | 41328-a.UCA | 10 | 0.47 | 6844 |
| | | V033-a.UCA | 10 | 0.03 | 2000 |
| | | V031-a.UCA | 10 | 0.71 | 6583 |
| | | 42056-b.UCA | 10 | 0.02 | 1390 |
| | | 5695-b.UCA | 10 | 0.04 | 1139 |
| | | PCT64 LMCA | 10 | 0.30 | 5512 |
| | | CH01 iGL | 0.477 | 0.13 | 4929 |
| | | CAP256-VRC26 UCA | 10 | 0.02 | 921 |
| | | PG9 iGL | 10 | 0.02 | 3176 |
| bnAbs | V2-Apex | 42056-a.01 | 0.00957 | 1.12 | 7482 |
| | | 40591-a.01 | 0.02247 | 0.87 | 10825 |
| | | RHA1.01 | 0.0046 | 1.02 | 11859 |
| | | T646-a.01 | 0.0046 | 1.27 | 8932 |
| | | 41328-a.01 | 0.0046 | 1.09 | 10689 |
| | | V033-a.01 | 0.00504 | 1.07 | 4250 |
| | | V031-a.01 | 0.0046 | 1.08 | 2151 |
| | | 42056-b.01 | 10 | 0.03 | 995 |
| | | PG9 | 0.0046 | 1.11 | 11686 |
| | | PG16 | 0.0046 | 0.98 | 11965 |
| | | PGT145 | 1.101 | 1.24 | 9236 |
| | | PGDM1400 | 0.0046 | 0.81 | 11256 |
| | | PCT64.35S | 0.0046 | 0.61 | 8007 |
| | | CH01 | 0.00769 | 0.07 | 8877 |
| | | CAP256.09 | 0.04755 | 0.52 | 8521 |
| | | CAP256.25 | 0.0046 | 0.76 | 9382 |
| | V3 Glycan | PGT121 | 0.0122 | 0.76 | 22496 |
| | | PGT128 | 0.00578 | 0.99 | 26611 |
| | | PGT130 | 0.05691 | 0.25 | 8821 |
| | | 10-1074 | 0.0046 | 0.92 | 22536 |
| | | PGT135 | 0.1166 | 0.20 | 14508 |
| | | 2G12 | 10 | 0.72 | 18989 |
| | CD4bs | VRC01 | 0.1242 | 0.37 | 13147 |
| | | VRC06 | 0.3822 | 0.28 | 15505 |
| | | HJ16 | 10 | 0.02 | 1460 |
| | | CH103 | 3.973 | 0.19 | 4684 |
| | | 3BNC60 | 0.01939 | 0.77 | 16274 |
| | gp120-41 | PGT151 | 0.0046 | 0.33 | 6637 |
| | | 35022 | 0.0046 | 0.37 | 11831 |
| | | VRC34.01 | 0.2124 | 0.86 | 2215 |
| nnAbs | V2i | 2909 | 6.015 | 0.07 | 1033 |
| | | 830A | 10 | 0.04 | 1398 |
| | | 697 | 10 | 0.01 | 1300 |
| | V3 nnAb | 2557 | 10 | 0.92 | 2464 |
| | | 3074 | 10 | 0.98 | 3041 |
| | CD4bs nnAb | b6 | 10 | 0.28 | 2785 |
| | | b12 | 10 | 0.11 | 1069 |
| | | F105 | 10 | 0.13 | 2340 |
| | CD4i | A32 | 10 | 0.04 | 1219 |
| | | 17b | 10 | 0.03 | 9020 |
| | | E51 | 10 | 0.05 | 974 |

B

C

D

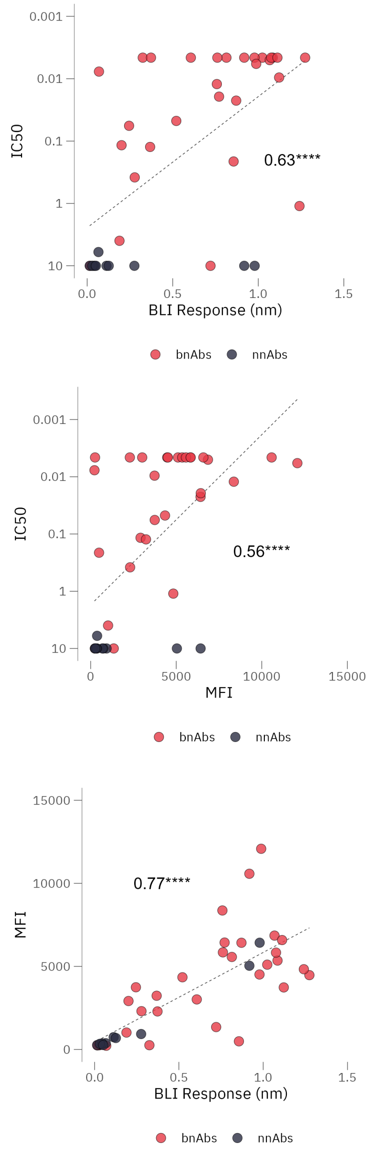

**Supplementary Figure 3. Further characterization of Q23-SCT27 antigenicity and expression, related to Figure 1.**

(A) Extended antigenicity data for Q23-SCT27 against a larger panel of bnAbs, non-nAbs and V2 apex UCAs (Human and Rhesus). Neutralization was tested starting at a concentration of 10 µg/ml and IC50 value of 10 denotes 50% neutralization was not reached at maximum concentration tested. For BLI, Q23-SCT27 trimer was tested at 500 nm and antibodies were tested at 10 µg/ml. For cell surface binding, Q23-SCT27 was appended to the wildtype TM from Q23 and tested for binding using flow cytometry.
(B–C) Correlation plots for neutralization versus soluble and cell surface expression, and (D) Soluble versus cell surface expression show significant positive correlation (Pearson's coefficient > 0.5). ****$P$< 0.0001.

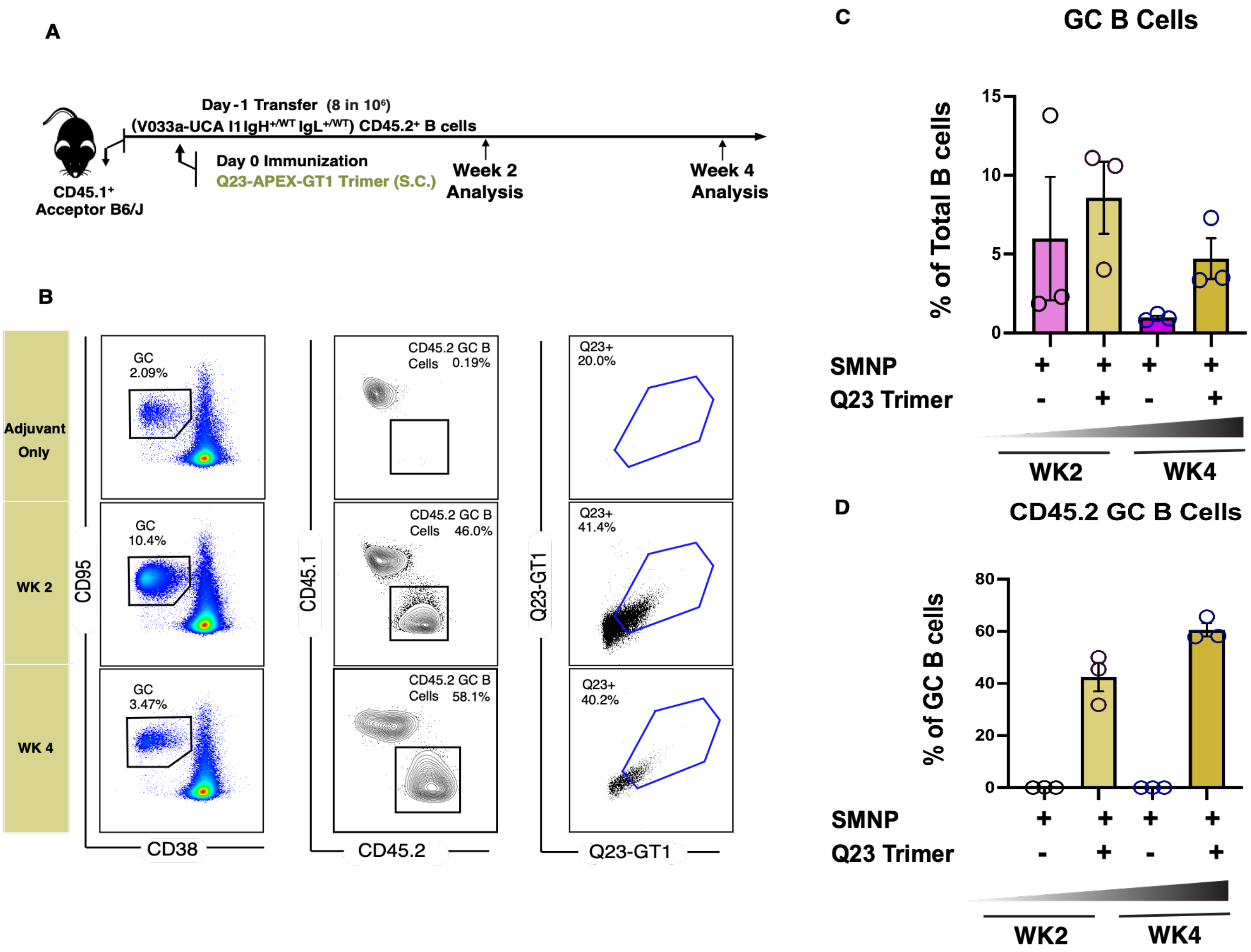
A
Day-1 Transfer (8 in 10⁶)
(V033a-UCA I1 IgH+/WT IgL+/WT) CD45.2+ B cells
Day 0 Immunization
Q23-APEX-GT1 Trimer (S.C.)
Week 2 Analysis
Week 4 Analysis
CD45.1+ Acceptor B6/J
B
Adjuvant Only
WK 2
WK 4
GC 2.09%
GC 10.4%
GC 3.47%
CD95
CD38
CD45.2 GC B Cells 0.19%
CD45.2 GC B Cells 46.0%
CD45.2 GC B Cells 58.1%
CD45.1
CD45.2
Q23+ 20.0%
Q23+ 41.4%
Q23+ 40.2%
Q23-GT1
C
GC B Cells
% of Total B cells
SMNP
Q23 Trimer
WK2
WK4
D
CD45.2 GC B Cells
% of GC B cells
SMNP
Q23 Trimer
WK2
WK4

**Supplementary Figure 4. Immunization with Q23-SCT after higher frequency of adoptive transfer leads to recruitment and activation of V033a-UCA I1 B cells, related to Figure 2.**

(A) Schematic of mouse adoptive transfer and immunization experiments. Mice received V033a-UCA I1 B cells through intravenous transfer one day prior to immunization with Q23-APEX-GT1 trimer adjuvanted with SMNP immunization. SMNP-only immunization served as a control.

(B) Representative FACS plots showing germinal centers, CD45.2 B cells in GCs and their binding to Q23-APEX-GT1 during weeks 2 and 4 post-immunization with Q23-APEX-GT1 trimer with SMNP.

(C, D) Quantification of GC B cells (C) and CD45.2 B cells in GCs (D) weeks 2 and 4 post-immunization.

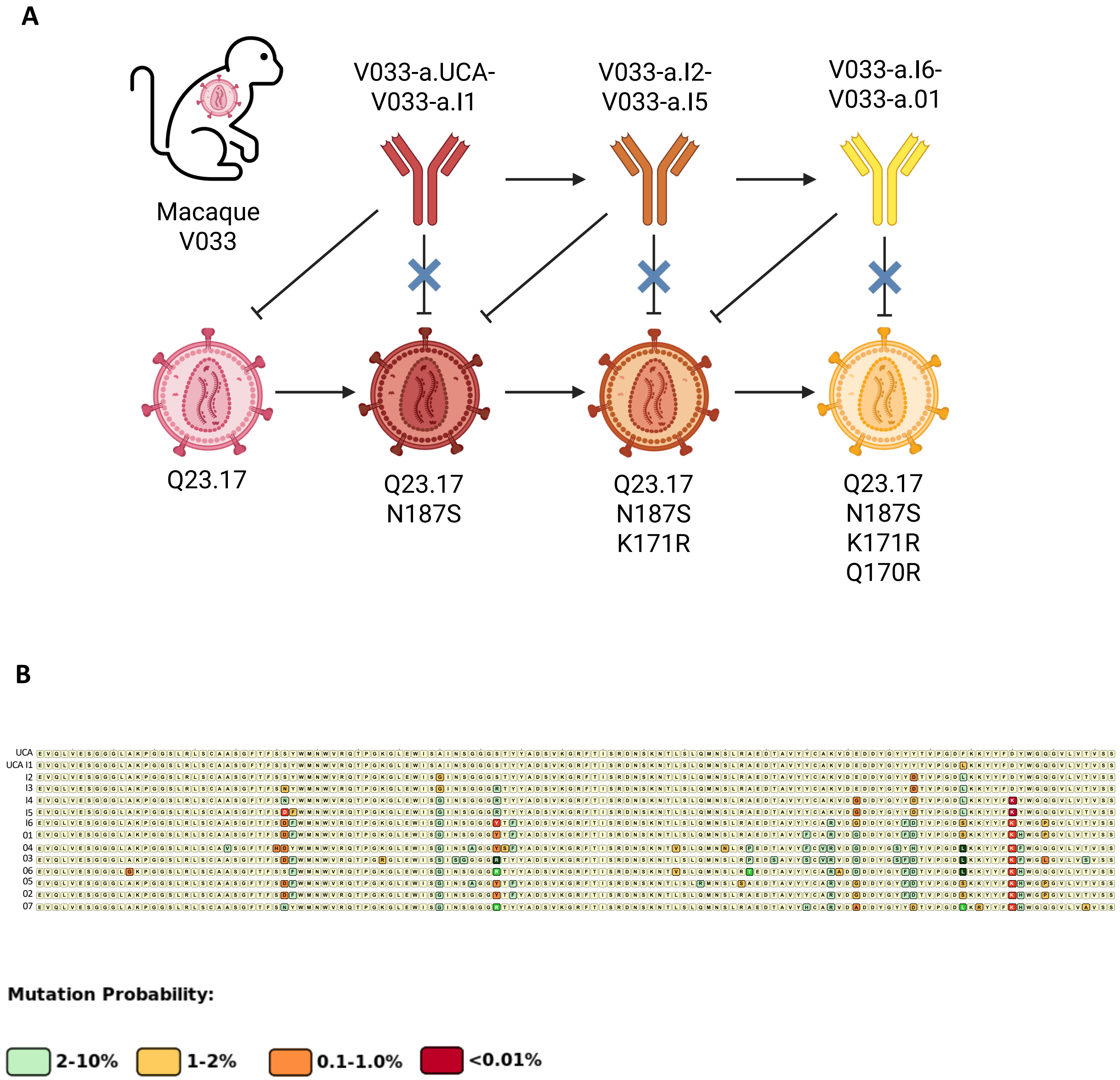
A
Macaque V033
V033-a.UCA-V033-a.I1
V033-a.I2-V033-a.I5
V033-a.I6-V033-a.01
Q23.17
Q23.17 N187S
Q23.17 N187S K171R
Q23.17 N187S K171R Q170R
B
UCA
UCA I1
I2
I3
I4
I5
I6
01
04
03
06
05
02
07
Mutation Probability:
2-10%
1-2%
0.1-1.0%
<0.01%

**Supplementary Figure 5. Macaque antibody features, related to Figure 4.**

(A) Schematic of SHIV infection and sequence isolation timeline.

(B) Probability of rare mutations as predicted by ARMaDILLO in mature macaque antibodies.

A

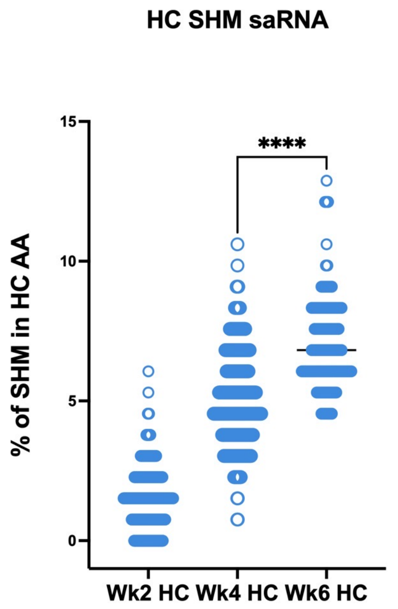


B

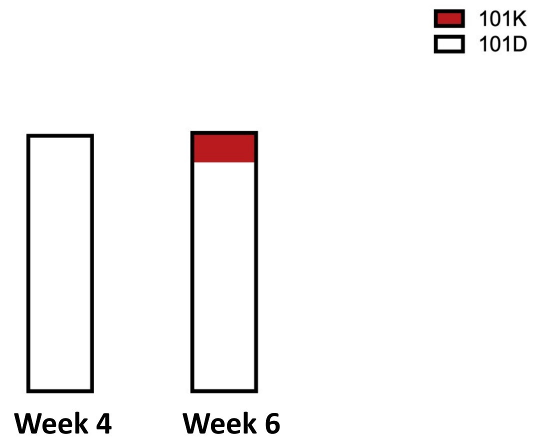


C

| Position | Mutation | Mutation probablity by ARMADiLLO | Frequency (%) at week 6 |
|---|---|---|---|
| HCDR1 | S31N | 1-2% | 99.36 |
| HCDR1 | S31D | 0.1-1% | 0.64 |
| HCDR2 | S56Y | 0.1-1% | 5.73 |
| HCDR2 | T57R | 1-2% | 37.58 |
| HCDR3 | E97D | 2-10% | 27.39 |
| HCDR3 | $Y100_CF$ | 2-10% | 34.39 |
| HCDR3 | $Y100_DD$ | 2-10% | 47.13 |
| HCDR3 | D100K | <0.01% | 11.46 |

**Supplementary Figure 6. Extended SHM analysis, related to Figure 3.**

(A) Total amino acid (AA) mutations in V033a-UCA I1 IGHV at weeks 2, 4, 6 post Q23-APEX-GT1 saRNA LNP immunization.

(B) Presence of rare Lysine mutation in HCDR3 post priming immunization with Q23-APEX-GT1.

(C) Frequency of selected mutations in V033a-UCA I1 after single priming immunization at week 6.

**A**

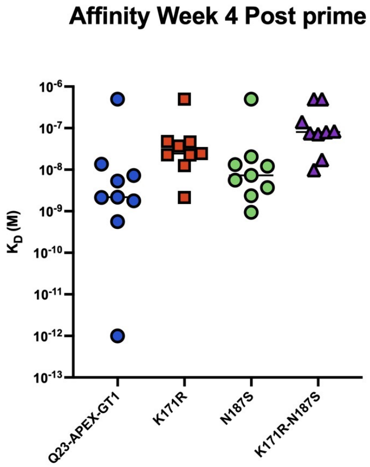


**B**

| | mAb1 | mAb2 | mAb3 | mAb4 | mAb5 | mAb6 | mAb7 | mAb8 | mAb9 | RMV033 P2_A6 |
|---|---|---|---|---|---|---|---|---|---|---|
| 246F3 | >50 | >50 | >50 | 15 | >50 | >50 | >50 | >50 | >50 | 0.034 |
| 25710 | 19 | 34 | >50 | 1.4 | >50 | 17 | 21 | >50 | >50 | 0.029 |
| BJOX0020000 | 5.8 | 4.8 | 12 | 0.662 | 17 | 5.5 | 7.2 | >50 | >50 | 0.040 |
| BG505.T332N | 3.6 | 4.4 | 7.0 | 0.766 | 41 | 2.1 | 2.6 | 49 | >50 | 0.020 |
| BG505.T332N.N160K | 6.5 | 6.7 | 10 | 1.2 | >50 | 3.0 | 3.4 | 41 | >50 | 0.321 |
| WITO.33 | >50 | >50 | >50 | 19 | >50 | 11 | 50 | >50 | >50 | 0.006 |
| Q23.17 | 0.141 | 0.062 | 0.209 | 0.029 | 0.560 | 0.053 | 0.096 | 0.766 | 5.2 | 0.017 |
| Q23.17 N160K | 2.4 | 1.2 | 25 | 0.268 | 4.1 | 0.398 | 0.699 | 3.8 | >50 | 0.002 |
| Q23.17 N187S | 1.7 | 1.4 | 5.1 | 0.269 | 26 | 1.3 | 1.3 | >50 | 37 | 0.116 |

mAb IC50 (ug/mL)

| <0.1 | 0.1-1.0 | 1.0-10 | 10-50 |
|---|---|---|---|

**C**

| Panel | Virus | mAb2 | mAb4 | mAb7 | V033-a.01 |
|---|---|---|---|---|---|
| Global panel | 246F3 | >50 | 15 | >50 | 0.034 |
| Global panel | X1632 | >50 | >50 | >50 | >50 |
| Global panel | 25710 | 34 | 1.4 | 21 | 0.029 |
| Global panel | CNE55 | >50 | >50 | >50 | >50 |
| Global panel | X2278 | >50 | >50 | >50 | 0.73 |
| Global panel | BJOX0020000 | 4.8 | 0.66 | 7.2 | 0.040 |
| Global panel | CNE8 | >50 | >50 | >50 | >50 |
| Global panel | CE0217 | >50 | >50 | >50 | 0.014 |
| Global panel | TRO.11 | >50 | >50 | >50 | >50 |
| Global panel | CE1176.A3 | >50 | >50 | >50 | >50 |
| Global panel | CH119.10 | >50 | 47 | >50 | 0.11 |
| | BG505.T332N | 4.4 | 0.77 | 2.6 | 0.020 |
| | BG505.T332N.N160K | 6.7 | 1.2 | 3.4 | 0.32 |
| | CAP256SU | >50 | >50 | >50 | >50 |
| | ZM233.6 | >50 | >50 | >50 | >50 |
| | WITO.33 | >50 | 19 | 50 | 0.006 |
| | T250-4 | >50 | >50 | >50 | 0.23 |
| | T250-4 N160K | >50 | >50 | >50 | >50 |
| | Q23.17 | 0.062 | 0.029 | 0.096 | 0.017 |
| | Q23.17 N160K | 1.2 | 0.27 | 0.70 | 0.002 |
| | Q23.17 N187S | 1.4 | 0.27 | 1.3 | 0.12 |
| | CH505 TF | >50 | >50 | >50 | >50 |
| | MT145K (SIVcpz) | >50 | >50 | >50 | 0.041 |
| | CAM13K (SIVcpz) | >50 | >50 | 30 | 0.045 |
| | C1080 | 0.25 | 0.072 | 0.23 | 0.006 |
| Hraber Panel | 9004SS_A3_4 | >50 | >50 | >50 | >50 |
| Hraber Panel | Q842.D12 | 18 | 0.82 | 8.5 | 0.027 |
| Hraber Panel | 249M_B10 | 44 | 4.8 | 43 | >50 |
| Hraber Panel | 7060101641A7(REV-) | >50 | >50 | >50 | >50 |
| Hraber Panel | HIV-001428-2.42 | 0.50 | 0.24 | 0.53 | 0.049 |
| Hraber Panel | HIV-16055-2.3 | 3.4 | 0.57 | 3.5 | 0.080 |
| Hraber Panel | HIV-25925-2.22 | >50 | 21 | >50 | 4.6 |
| Hraber Panel | 3728.v2.c6 | >50 | >50 | >50 | >50 |
| Hraber Panel | RPW-0510.2 | >50 | >50 | >50 | >50 |
| Hraber Panel | H029.12 | >50 | >50 | >50 | >50 |
| Hraber Panel | SPK-0525.13 | >50 | 18 | 15 | 0.11 |

**Supplementary Figure 7. Affinity, neutralization breadth of prime derived antibodies, related to Figure 4.**

(A) BLI affinity values of antibodies derived 6 weeks post-prime against autologous Q23 Env and escape variant Envelopes.

(B) Neutralization breadth and potency of selected antibodies from 4 weeks post-prime and macaque mature V033 (RM V033 P2A6) against autologous (Q23), escape variants (Q23.N187S, Q23.K171R.N187S) and limited panel of 6 heterologous tier-2 HIV-1 strains. Antibodies from Trimer+SMNP immunization are marked in magenta and mAbs from Q23-APEX-GT1 saRNA LNP are marked in blue.

(C) IC50 of selected three antibodies against a 37-member panel of HIV-1 strains including 11 from Tier-2 Global panel and 11 from Hraber panel.

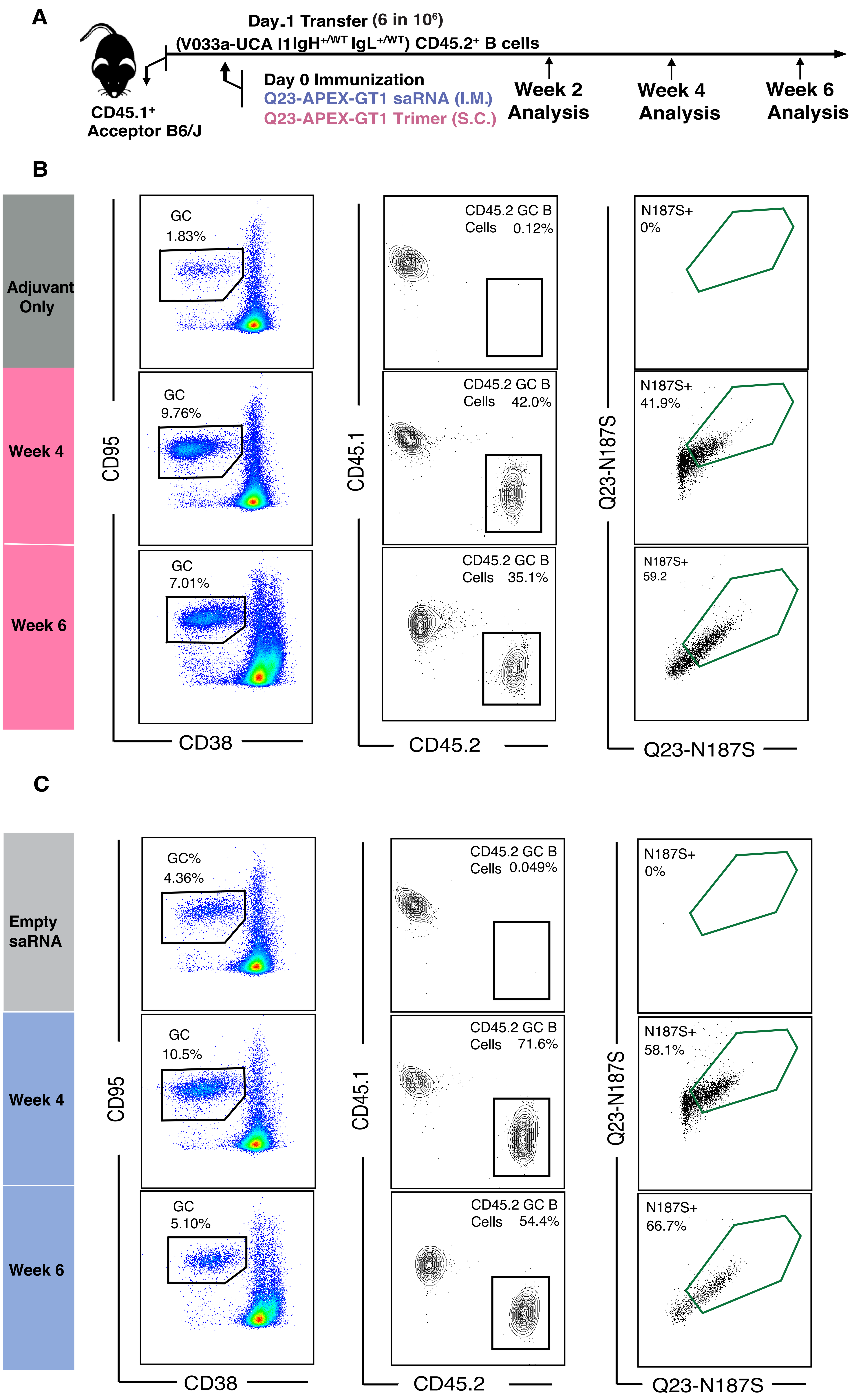
A
Day-1 Transfer (6 in 10⁶)
(V033a-UCA I1IgH+/WT IgL+/WT) CD45.2+ B cells
CD45.1+
Acceptor B6/J
Day 0 Immunization
Q23-APEX-GT1 saRNA (I.M.)
Q23-APEX-GT1 Trimer (S.C.)
Week 2
Analysis
Week 4
Analysis
Week 6
Analysis
B
Adjuvant Only
Week 4
Week 6
GC
1.83%
GC
9.76%
GC
7.01%
CD45.2 GC B Cells 0.12%
CD45.2 GC B Cells 42.0%
CD45.2 GC B Cells 35.1%
N187S+
0%
N187S+
41.9%
N187S+
59.2
CD95
CD38
CD45.1
CD45.2
Q23-N187S
Q23-N187S
C
Empty saRNA
Week 4
Week 6
GC%
4.36%
GC
10.5%
GC
5.10%
CD45.2 GC B Cells 0.049%
CD45.2 GC B Cells 71.6%
CD45.2 GC B Cells 54.4%
N187S+
0%
N187S+
58.1%
N187S+
66.7%
CD95
CD38
CD45.1
CD45.2
Q23-N187S
Q23-N187S

**Supplementary Figure 8. Q23-APEX-GT1 primed V033a-UCA I1 B cells can bind Q23 escape variant Q23-N187S, related to Figures 3 and 7.**

(A) Schematic presentation of mouse adoptive transfer and immunization experiments. SMNP adjuvant without trimer is used as control for protein immunizations and LNPs containing an unrelated saRNA was used as control for Q23-APEX-GT1 saRNA LNP immunization.

(B, C) Representative FACS plots showing germinal centers, CD45.2 B cells in GCs and their binding to the escape variant Q23-N187S during weeks 4 and 6 post immunization with Q23-APEX-GT1 trimer with SMNP (B) or LNPs containing Q23-APEX-GT1 saRNA (C).

**A**

| | Empty saRNA | | | Q23-APEX-GT1 saRNA | | | SMNP Only | | | Q23-APEX-GT1 +SMNP | | |
|---|---|---|---|---|---|---|---|---|---|---|---|---|
| **MULV** | <100 | <100 | <100 | <100 | <100 | <100 | <100 | <100 | <100 | <100 | <100 | <100 |
| **Q23.17 WT** | <100 | <100 | <100 | <100 | <100 | <100 | <100 | <100 | <100 | 238 | 151 | 465 |
| **Q23.17 N160K** | <100 | <100 | <100 | <100 | <100 | <100 | <100 | <100 | <100 | <100 | <100 | <100 |

| **1/ID50** | 100 - 1000 | 1000 - 10000 | >10000 |
|---|---|---|---|

**B**

| | Empty saRNA | | | Q23-APEX-GT1 saRNA | | | SMNP Only | | | Q23-APEX-GT1 +SMNP | |
|---|---|---|---|---|---|---|---|---|---|---|---|
| | 2R | 1L1R | | 2R | 1L1R | | 2R | 1L1R | | 2R | 1L1R |
| **Q23.17 WT** | <100 | <100 | | <100 | <100 | | <100 | <100 | | 109 | 115 |
| **Q23.17 N160K** | <100 | <100 | | <100 | <100 | | <100 | <100 | | <100 | <100 |

| **1/ID50** | 100 - 1000 | 1000 - 10000 | >10000 |
|---|---|---|---|

**Supplementary Figure 9. Priming leads to minimal serum neutralization, related to Figure 7.**

Reciprocal serum $ID_{50}$ values of animals receiving Q23-APEX-GT1 Prime at week 4 (A) and week 6 (B) post-immunization against Q23.17, Q23.17 N160K.

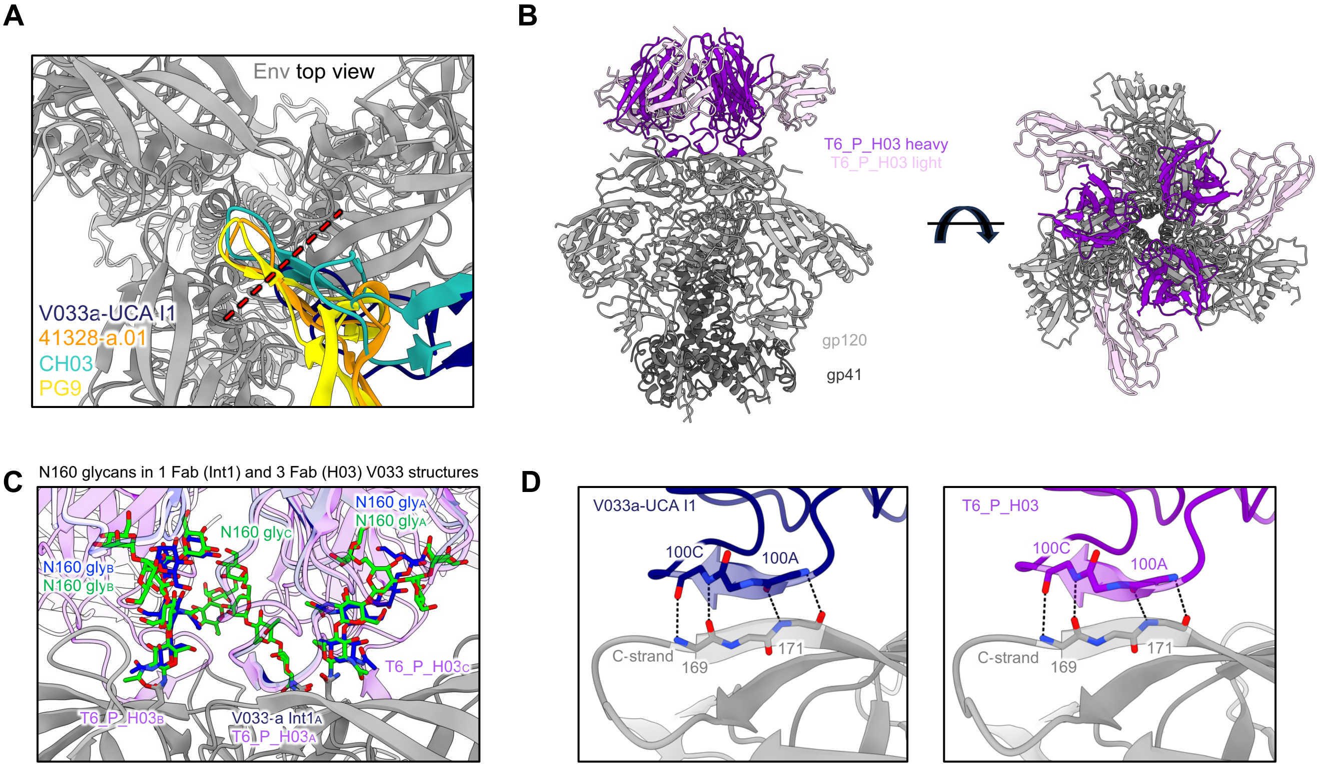

A
Env top view
V033a-UCA I1
41328-a.01
CH03
PG9
B
T6_P_H03 heavy
T6_P_H03 light
gp120
gp41
C
N160 glycans in 1 Fab (Int1) and 3 Fab (H03) V033 structures
N160 gly$_A$
N160 gly$_A$
N160 gly$_C$
N160 gly$_B$
N160 gly$_B$
T6_P_H03$_C$
T6_P_H03$_B$
V033-a Int1$_A$
T6_P_H03$_A$
D
V033a-UCA I1
100C
100A
C-strand
169
171
T6_P_H03
100C
100A
C-strand
169
171

**Supplementary Figure 10. Select comparative structural features of V033a-UCA I1 and prime-derived antibody T6_P_H03**

(A) Top envelope trimer view for the gp120 alignment of V033-a I1 (PDB-XXX) and axe-like V2 apex bnAbs 41328-a.01 (PDB-9BTL), CH03 (PDB-5ESZ), and PG9 (PDB-7T77). Only the Fab HCDR3 from each respective complex structure is shown for clarity. The red dashed line denotes the edge of the V033a UCA I1 HCDR3 tip to highlight the contrast in its relative positioning compared to other axe-like lineages, the latter of which all intersect with the trimer 3-fold axis.

(B) Orthogonal views for the cryo-EM structure of T6_P_H03 in complex with Q23-APEX-GT1 envelope trimer. The top view (right) reveals the middle of the trimer to be unencumbered by Fab HCDR3, similar to V033a-UCA I1, thereby allowing 3 Fabs to bind at the V2 apex.

(C) Comparing the conformation of N160 glycans recognized by 1 Fab bound (V033-a I1) and 3 Fab bound (T6_P_H03) envelope complexes from V033-a variant cryo-EM structures. The structures of V033a UCA I1 and T6_P_H03 are superimposed by gp120 alignment of protomer A. The conformation of the two N160 glycans recognized by a single V033a UCA I1 Fab (blue glycans) is compatible with recognition by multiple Fabs (green glycans).

(D) V033a-UCA I1 and T6_P_H03 HCDR3s each recognize the C-strand with four identical antiparallel mainchain hydrogen bonds.

A

**Q23-APEX-GT1 Prime**
**Q23 N187S Boost**

| | m1 | m2 | m3 | m4 |
|---|---|---|---|---|
| Q23.17 WT | 1039 | 487 | 3378 | 2221 |
| Q23.17 N160K | 198 | <100 | 3574 | 979 |
| Q23.17 N187S | <100 | <100 | 184 | 293 |

B

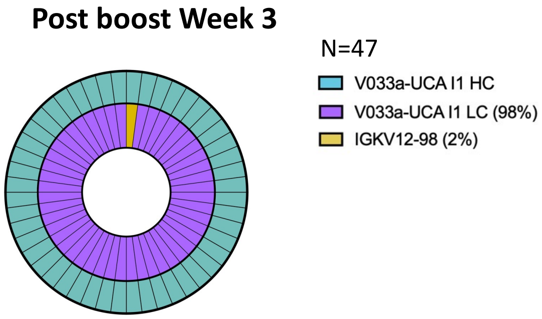


C

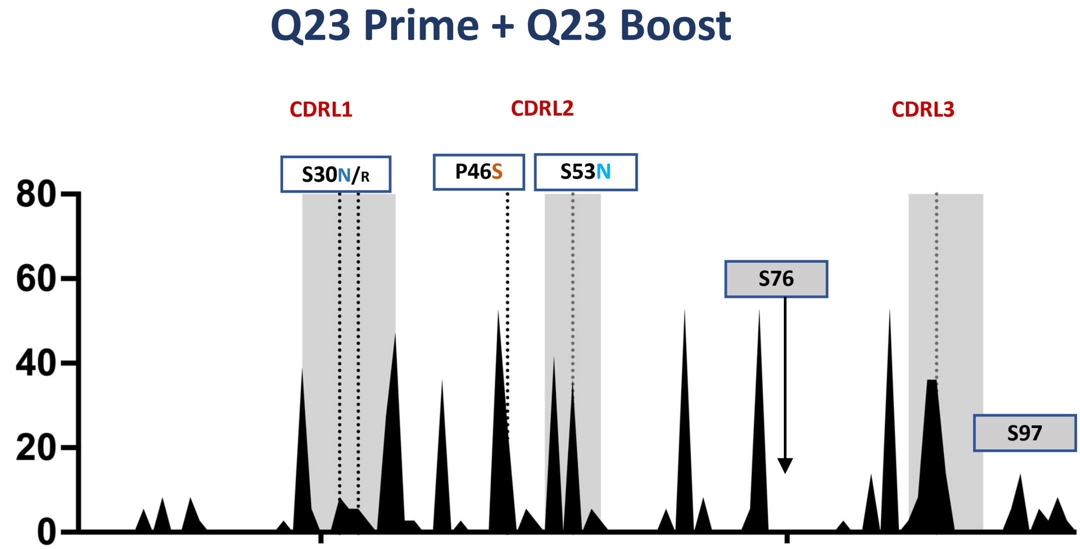


D

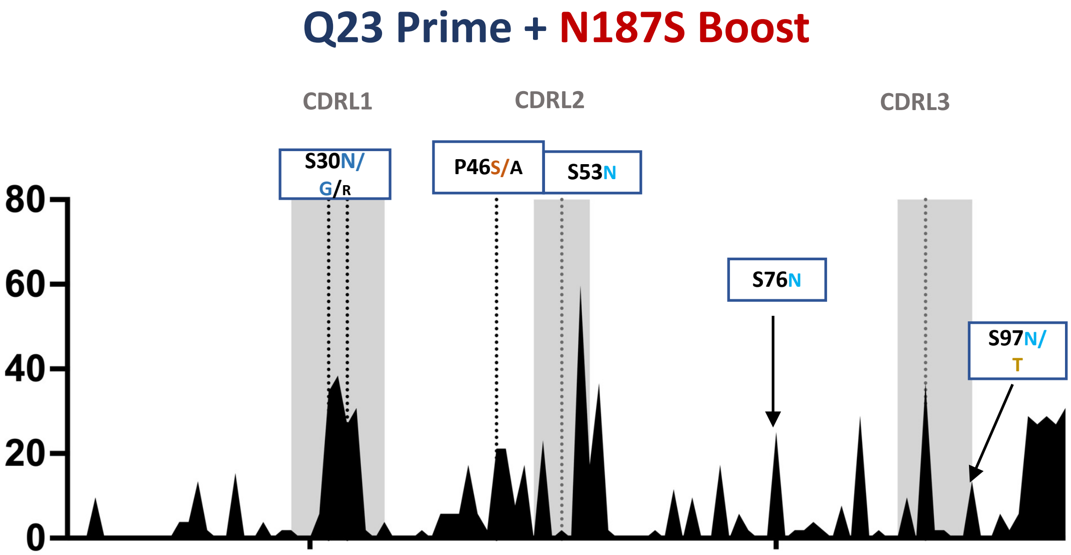

**Supplementary Figure 11. Heterologous boost leads to expanded serum neutralization and LC usage in homologous vs N187S boost, related to Figures 6 and 7.**

(A) Reverse ID50 of serum of animals receiving Q23-GT1 Prime and Q23-N187S boost against Q23.17, Q23.17 N160K and Q23.17 N187S escape variant.

(B) Light chain usage after boost.

(C,D) LC mutation frequencies after week 3 post immunization of V033_UCA I1 KI LC after Q23 boost (C) and N187S Boost (D). Some of the selected mutations present in mature V033 lineage bnAbs is represented in Red and intermediate mutations are represented in blue.

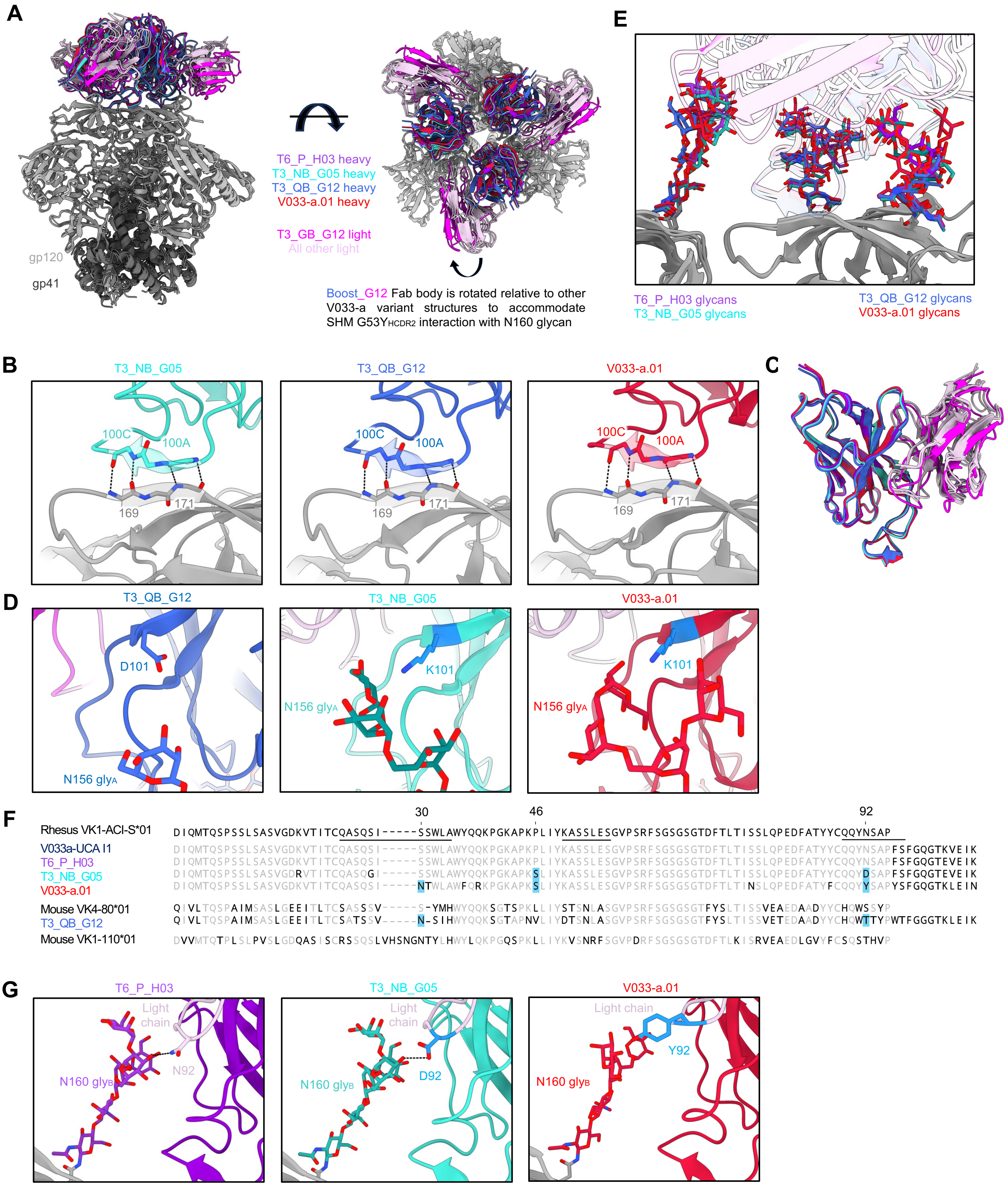
A
T6_P_H03 heavy
T3_NB_G05 heavy
T3_QB_G12 heavy
V033-a.01 heavy
T3_GB_G12 light
All other light
gp120
gp41
Boost_G12 Fab body is rotated relative to other V033-a variant structures to accommodate SHM G53Y HCDR2 interaction with N160 glycan
E
T6_P_H03 glycans
T3_NB_G05 glycans
T3_QB_G12 glycans
V033-a.01 glycans
B
T3_NB_G05
T3_QB_G12
V033-a.01
100C
100A
169
171
C
D
T3_QB_G12
T3_NB_G05
V033-a.01
D101
K101
N156 glyA
F
30
46
92
Rhesus VK1-ACI-S*01 DIQMTQSPSSLSASVGDKVTITCQASQSI-----SSWLAWYQQKPGKAPKPLIYKASSLESGVPSRFSGSGSGTDFTLTISSLQPEDFATYYCQQYNSAP
V033a-UCA I1 DIQMTQSPSSLSASVGDKVTITCQASQSI-----SSWLAWYQQKPGKAPKPLIYKASSLESGVPSRFSGSGSGTDFTLTISSLQPEDFATYYCQQYNSAPFSFGQGTKVEIK
T6_P_H03 DIQMTQSPSSLSASVGDKVTITCQASQSI-----SSWLAWYQQKPGKAPKPLIYKASSLESGVPSRFSGSGSGTDFTLTISSLQPEDFATYYCQQYNSAPFSFGQGTEVEIK
T3_NB_G05 DIQMTQSPSSLSASVGDRVTITCQASQGI-----SSWLAWYQQKPGKAPKSLIYKASSLESGVPSRFSGSGSGTDFTLTISSLQPEDFATYYCQQYDSAPFSFGQGTEVEIK
V033-a.01 DIQMTQSPSSLSASVGDKVTITCQASQSI-----NTWLAWFQRKPGKAPKSLIYKASSLESGVPSRFSGSGSGTDFTLTINSLQPEDFATYFCQQYYSAPYSFGQGTKLEIN
Mouse VK4-80*01 QIVLTQSPAIMSASLGEEITLTCSASSSV------S-YMHWYQQKSGTSPKLLIYSTSNLASGVPSRFSGSGSGTFYSLTISSVEAEDAADYYCHQWSSYP
T3_QB_G12 QIVLTQSPAIMSASLGEEITLTCSATSSV------N-SIHWYQQKSGTAPNVLIYDTSNLASGVPSRFSGSGSGTFYSLTISSVETEDAADYYCHQWTTYPWTFGGGTKLEIK
Mouse VK1-110*01 DVVMTQTPLSLPVSLGDQASISCRSSQSLVHSNGNTYLHWYLQKPGQSPKLLIYKVSNRFSGVPDRFSGSGSGTDFTLKISRVEAEDLGVYFCSQSTHVP
G
T6_P_H03
T3_NB_G05
V033-a.01
Light chain
N92
D92
Y92
N160 glyB

**Supplementary Figure 12. Select comparative structural features of V033-a lineage variants.**

(A) Orthogonal views for the gp120 alignment of T6_P_H03, T3_NB_G05, T3_QB_G12, and V033-a.01 cryo-EM structures in complex with Env SOSIP trimers.

(B) Boost-derived antibodies each recognize the C-strand with four antiparallel mainchain hydrogen bonds identical to rhesus V033-a lineage members, including V033-a.01.

(C) Superimposition of T6_P_H03, T3_NB_G05, T3_QB_G12, and V033-a.01 Fabs from their respective complex structures. The use of a mouse-derived light chain does not alter the structure of antibody T3_QB_G12 and likely does not contribute to the unique angle of approach relative to all other V033-a lineage variants. Heavy and light chains are colored similarly to panel A.

(D) The improbable HCDR3 K101 mutation in V033-a lineage variants T3_NB_G05 and V033-a.01 mediates recognition of N156 glycan from protomer A. In the T3_QB_G12 structure bearing the germline residue D101, the visible reconstruction density reveals N156 glycan on protomer A to not extend as far and does not interact with this residue.

(E) Comparing the conformation of apical glycans recognized by T6_P_H03, T3_NB_G05, T3_QB_G12, and V033-a.01. A single Fab recognizing the C-strand of protomer A binds to the N160 and N156 glycans from protomer A and N160 glycan from protomer B. All apical glycans adopt highly similar conformations when recognized by V033-a lineage variants. Glycans are colored according to the corresponding antibody heavy chain from panel A. Fab structures are made transparent to better show glycans. Only one Fab per structure is shown for clarity.

(F) Light chain amino acid sequences of T6_P_H03, T3_NB_G05, T3_QB_G12, V033-a.01, and V033-a I1 are aligned to germline rhesus VK1-ACI-S*01 gene. Antibody T3_QB_G12 is derived from a mouse light chain, and its germline gene VK4-80*01 is included in the alignment as well to visualize T3_QB_G12 light chain SHM. Many V033-a variant mouse antibodies were also derived from lights chains expressing VK1-110*01, and this sequence is included in the alignment as well. Residues matching the reference are depicted in light gray and nonmatching residues are depicted in black. The LCDRs are overscored in the V033-a I1 sequence. Positions of somatic hypermutation shared by one or two murine antibodies with V033-a.01 are highlighted in light blue.

(G) V033-a lineage variant antibodies bearing rhesus light chains recognize N160 glycan from protomer B using LCDR3 residue 92. The D92 SHM in T3_NB_G05 does not significantly alter the N160 glycan-recognizing paratope over germline N92 present in T6_P_H03; however, the V033-a.01 LCDR3 adopts a slightly different conformation and uses aromatic stacking to recognize N160 glycan via Y92 SHM.

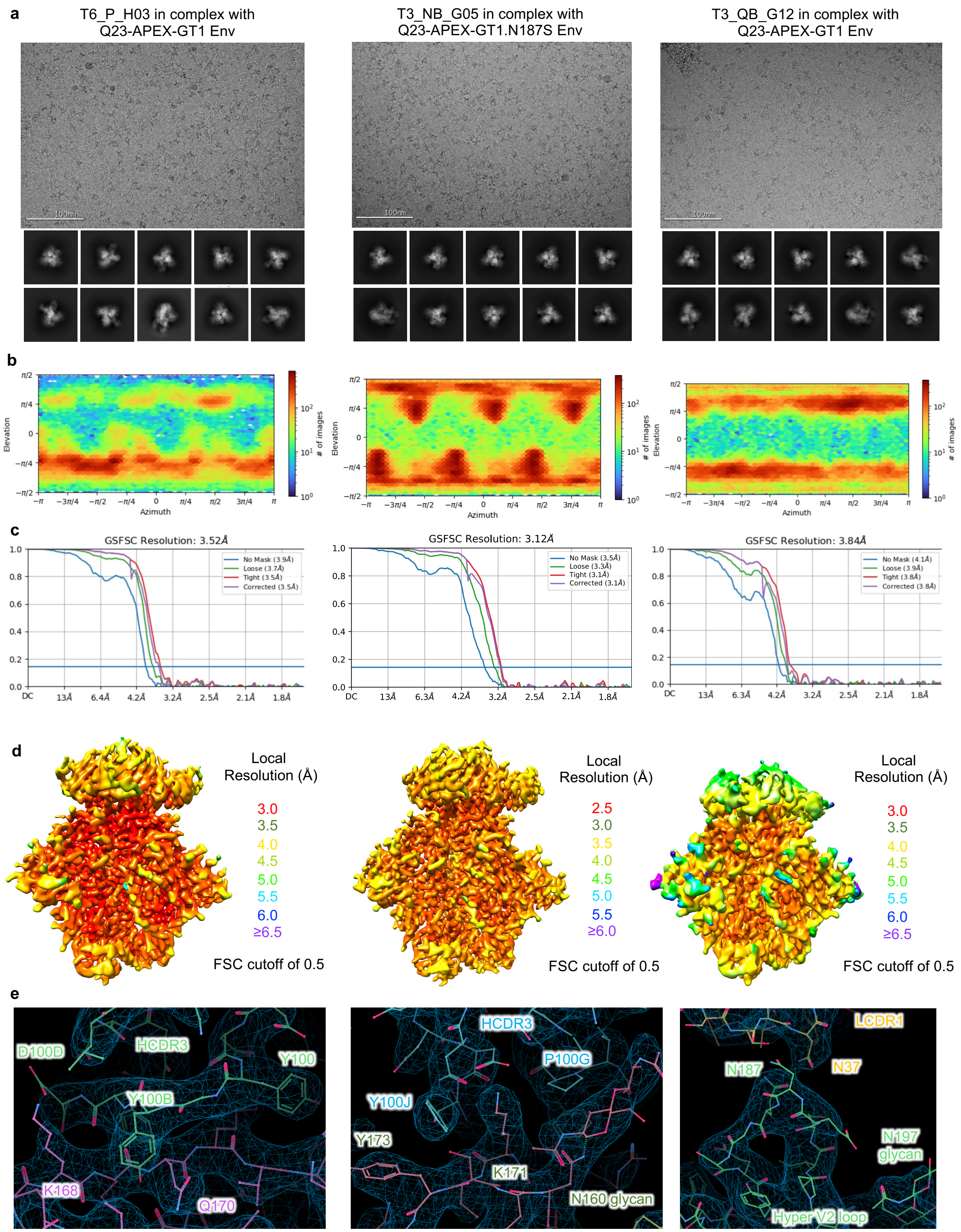
a
T6_P_H03 in complex with
Q23-APEX-GT1 Env
T3_NB_G05 in complex with
Q23-APEX-GT1.N187S Env
T3_QB_G12 in complex with
Q23-APEX-GT1 Env
100nm
b
Elevation
Azimuth
### of images
c
GSFSC Resolution: 3.52Å
No Mask (3.9Å)
Loose (3.7Å)
Tight (3.5Å)
Corrected (3.5Å)
GSFSC Resolution: 3.12Å
No Mask (3.5Å)
Loose (3.3Å)
Tight (3.1Å)
Corrected (3.1Å)
GSFSC Resolution: 3.84Å
No Mask (4.1Å)
Loose (3.9Å)
Tight (3.8Å)
Corrected (3.8Å)
d
Local
Resolution (Å)
3.0
3.5
4.0
4.5
5.0
5.5
6.0
≥6.5
FSC cutoff of 0.5
Local
Resolution (Å)
2.5
3.0
3.5
4.0
4.5
5.0
5.5
≥6.0
FSC cutoff of 0.5
Local
Resolution (Å)
3.0
3.5
4.0
4.5
5.0
5.5
6.0
≥6.5
FSC cutoff of 0.5
e
D100D
HCDR3
Y100
Y100B
K168
Q170
HCDR3
P100G
Y100J
Y173
K171
N160 glycan
LCDR1
N187
N37
N197
glycan
Hyper V2 loop

**Supplementary Figure 13. Single-particle cryo-EM validation for murine V033-a antibodies in complex with HIV envelope.**

(A) Representative raw micrograph with representative 2D class averages of picked particles shown below.

(B) Orientations of all particles used in the final refinement are shown as a heatmap.

(C) Gold-standard fourier shell correlation (FSC) curves with auto-tightening using a non-uniform refinement with C3 symmetry.

(D) Local resolution estimation of the full map is shown as generated through cryoSPARC using an FSC cutoff of 0.5.

(E) Example of high-resolution cryo-EM 3D reconstruction density to highlight regions of the Fab:trimer interface.

### Supplementary Table 1 – cryoEM statistics

**Supplementary Table 1 | Cryo-EM data collection, processing, and refinement validation statistics**

| | T6_P_H03 in complex with Q23-APEX-GT1 Env | T3_NB_G05 in complex with Q23-APEX-GT1.N187S Env | T3_QB_G12 in complex with Q23-APEX-GT1 Env |
|---|---|---|---|
| **PDB** | **9OOM** | **9OOG** | **9OOK** |
| **EMDB** | **70666** | **70663** | **70664** |
| **Data collection & processing** | | | |
| Microscope | FEI Titan Krios | FEI Titan Krios | FEI Titan Krios |
| Camera | Gatan K3 | Gatan K3 | Gatan K3 |
| Magnification | 105,000x | 105,000x | 105,000x |
| Voltage (kV) | 300 | 300 | 300 |
| Electron dose ($e^-/Å^2$) | 58 | 58 | 58 |
| Defocus range (μm) | 0.8 - 2.0 | 0.8 - 2.0 | 1.0 - 2.5 |
| Pixel size (Å) | 0.83 | 0.83 | 0.83 |
| Micrographs collected | 4,956 | 8,638 | 8,970 |
| Software | cryoSPARC v4.1 | cryoSPARC v4.1 | cryoSPARC v4.1 |
| Micrographs used | 4,183 | 7,862 | 7,991 |
| Refined particles | 140,657 | 211,774 | 140,366 |
| Symmetry imposed | C3 | C3 | C3 |
| Map Resolution (Å) | 3.52 | 3.12 | 3.84 |
| FSC threshold | 0.143 | 0.143 | 0.143 |
| **Refinement & validation** | | | |
| Initial model used | 9BNP | 9BNP | 9BNP |
| Software | Phenix 1.21 | Phenix 1.21 | Phenix 1.21 |
| Number of residues | | | |
| Protein | 2,433 | 2,433 | 2,430 |
| Ligand | 145 | 141 | 132 |
| Map CC | 0.83 | 0.85 | 0.80 |
| R.m.s. deviations | | | |
| Bond lengths (Å) | 0.006 | 0.005 | 0.004 |
| Bond angles (˚) | 1.01 | 0.897 | 0.931 |
| EMRinger score | 3.01 | 2.72 | 1.78 |
| MolProbity score | 1.56 | 1.27 | 1.43 |
| Clashscore | 4.29 | 2.08 | 3.14 |
| Rotamer outliers (%) | 0 | 0 | 0 |
| Ramachandran plot | | | |
| Favored (%) | 95.0 | 95.9 | 95.3 |
| Allowed (%) | 5.0 | 4.1 | 4.7 |
| Outliers (%) | 0 | 0 | 0.0 |
